# *StemCNV-check*: a pipeline for human pluripotent stem cell (hPSC) genomic integrity control using SNP array data and copy number variant scoring

**DOI:** 10.64898/2025.12.01.691615

**Authors:** Nicolai von Kügelgen, Valeria Fernandez Vallone, Katarzyna A. Ludwik, Mariia Babaeva, Sebastian Diecke, Dieter Beule, Harald Stachelscheid

**Author notes:** both authors contributed equally (Nicolai von Kügelgen, Valeria Fernandez Vallone). senior last-authors (Dieter Beule, Harald Stachelscheid).

## Abstract

Human pluripotent stem cells (hPSCs) and other continuously cultured cell lines are prone to acquiring mutations and genomic aberrations over time, even when derived from well-characterized cell banks. To ensure experimental reproducibility and maintain cell line integrity, routine monitoring for genomic abnormalities is essential. Single nucleotide polymorphism (SNP) arrays represent a cost-effective and widely accessible method for detecting copy number variations (CNVs) with genome-wide resolution, making them particularly suitable for quality control (QC) in cell culture systems.

Despite the established utility of SNP arrays for CNV detection, there remains a lack of comprehensive, user-friendly software solutions that support end-to-end analysis tailored to hPSC line quality assessment. Existing tools are either limited to discrete analysis steps requiring specialized bioinformatics expertise or are proprietary solutions that do not adequately address the specific needs of cell line monitoring.

To bridge this gap, we developed an accessible and integrated analysis pipeline for SNP array-based QC of hPSC lines. The pipeline facilitates all stages of analysis—from raw data processing to the generation of interpretable reports—and includes specialized features such as sample-to-reference comparison, a CNV scoring system according to CNV biological impact, single nucleotide variation (SNV) evaluation and identity verification via SNP genotyping profiles, all tailored to hPSC. We benchmarked the pipeline against established methodologies and implemented strategies to enhance CNV detection reliability through expert-guided improvement process.

## Introduction

Human pluripotent stem cells (hPSC) are essential tools for basic and pre-clinical research, including disease modelling, target discovery, drug screening, and cell therapy. Maintaining their genetic integrity is critical for reproducible research and clinical safety, as emphasized in the recent ISSCR Standards for Human Stem Cell Use in Research ^1^ (https://www.isscr.org/guidelines).

Due to their self-renewal capacity, hPSC can proliferate indefinitely *in vitro*, but long-term culture often leads to genomic alterations, including large chromosomal changes, sub-chromosomal copy number variations (CNVs), and single nucleotide polymorphisms (SNPs) ^2^. Their rate of acquisition is influenced by culture conditions, such as medium, matrix, oxygen levels, passaging frequency, cell density, and dissociation methods. Moreover, CRISPR/Cas-mediated gene editing and subsequent single-cell cloning impose additional stress, further increasing the risk of genomic instability ^2^.

Recurrent genomic aberrations are frequently observed in hPSC cultures ^2–4^. Among the most observed CNVs are gains on chromosomes 1q, 12p, and 20q, as well as losses on 17p, which have been linked to increased self-renewal and proliferation capacities ^5^. These aberrations raise concerns about the functional and clinical implications of using genetically unstable hPSC lines. Moreover, regions such as 20q11.21 have been implicated in promoting clonal expansion under selective pressure, particularly during prolonged *in vitro* culture ^6^. Not only CNVs, in the form of regional gains or losses, but also SNVs in key genes are common abnormalities affecting hPSC proliferation and differentiation potential ^7–12^.

Detection of genomic aberrations requires methodologies matched to the type and scale of the variant. Classical G-banding karyotyping is effective for identifying whole-chromosome aneuploidies and large structural alterations, such as translocations, inversions, deletions, and duplications involving mega base-sized regions. However, smaller CNVs—often with functional impact—frequently escape detection by conventional cytogenetics and require more sensitive techniques, including SNP arrays, comparative genomic hybridization arrays, or whole-genome/exome sequencing (WGS/WES). These methods differ in sensitivity, specificity, turnaround time, and cost, factors that determine their suitability for hPSC quality control (QC), particularly during line derivation, genome editing, and biobanking. Among them, SNP arrays provide reliable resolution of submicroscopic CNVs with relatively fast turnaround (7–10 days) and low cost (∼€35 per sample, depending on array type).

Although many CNV calling algorithms and analysis tools exist, they rarely address the specific needs of hPSC research, such as integrated QC and context-specific interpretation ^13–15^. Crucially, no current solution provides a complete, stem cell–tailored workflow—from SNP array raw data processing to biologically meaningful interpretation—allowing researchers to assess CNVs in terms of their impact on hPSC function and downstream applications. Filling this gap would markedly improve data quality and reproducibility across laboratories.

We developed and validated *StemCNV-check*, a robust SNP-array analysis pipeline for CNV detection that surpasses commonly used tools in both accuracy and user-friendliness. Designed for routine use by stem cell biologists, it installs easily on Linux or Windows systems via Windows Subsystem for Linux (WSL). In addition to detecting CNV gains, losses, and loss of heterozygosity (LOH), it evaluates protein coding SNV detected by the array according to potential protein function alterations. All variant calls are fully annotated and can be instantly compared to a reference sample, enabling rapid validation of results.

To support interpretation, *StemCNV-check* introduces the *Check-Score*, ranking CNVs by size and biological relevance, and presents results in an interactive HTML report for intuitive genomic integrity assessment. It also enables hPSC line authentication and detection of sample misidentification and can assess on- and off-target effects of genome editing. Scalable from single samples to large cohorts, *StemCNV-check* enables reproducible processing by providing complete a standardized workflow that advances data quality and reporting in hPSC research. It is easy to install and update using bioconda. All code is open-source with a permissive license fulfilling FAIR4RS criteria ^16^, see section “Availibilty” for full details.

## Material and methods

### Data generation

A total of 381 hiPSC lines were analyzed, 328 from the Berlin Institute of Health Core Unit Pluripotent Stem Cells and Organoids (will be called BIH Charité dataset), 25 from the Max Delbrück Center (MDC), 12 from Fraunhofer-Institute for Translational Medicine and Pharmacology (ITMP) and 16 from Institute of Molecular Biotechnology (IMBA). Detailed metadata (passage number, culture conditions, and genetic modifications etc.) is provided in **Supplementary Table 1**. Genomic DNA was extracted and analyzed using the Infinium Global Screening Array v3.0 (GSAMD-24v3-0-EA_20034606; Illumina) and processed by Life & Brain GmbH (Bonn). For validation purposes, whole-genome sequencing was also performed on 13 selected hiPSC samples.

Whole-genome sequencing (WGS) was performed by the BIH Genomics Facility using the Illumina DNA PCR-Free Prep, Tagmentation kit (#20041795) and sequenced on a NovaSeq 6000 S4 flow cell (2×150 bp, ∼1000 million reads, ∼70× mean coverage. Reference data from six genomic DNA samples were obtained from the NIGMS Human Genetic Cell Repository (Coriell Institute) and genotyped using the SNP array platform (**Supplementary Table 1**). Corresponding WGS fastq files were downloaded from the Genome in a Bottle (GIAB) dataset via NCBI in July 2023 (**Supplementary Table 2**), ensuring consistent coverage and processing across all samples.

Additionally a reprogramming cohort dataset was used, which contained fibroblasts from 6 donors that were equally mixed in a pool and subjected to reprogramming using Sendai Virus as previously described ^17^. After colony picking, 30 hiPSC clones were expanded (passage 1-3). Genomic DNA was extracted from both parental fibroblasts and derived hiPSC clones. SNP array was performed as described above.

### Data processing

Array data were processed using both *GenomeStudio* software (Illumina) and our custom-developed tool, *StemCNV-check* (described below). *GenomeStudio* (v2.0.4) was used to analyze the data in two separate project instances: one for samples from our in-house hiPSC collection (used in Figure 1) and another for samples with matching WGS data (used in Figure 2). All *GenomeStudio* analyses employed default settings: Gen Call Threshold=0.15, no cluster recalculations, and CNV detection using the cnvPartition plugin (v3.2.0, default parameters). *StemCNV-check* was initially applied in a prototype version for the analyses presented in **Figure 1**, and later with the finalized release v1.0 for further analyses. Specific implementation details are provided in subsequent sections. WGS data were processed using the established in-house snappy pipeline (https://github.com/bihealth/snappy-pipeline). Briefly, reads were aligned to the human genome reference GRCh37 (hs37d5) using bwa-mem2 (v2.2.1; ^18^). CNV analysis was performed using GATK4’s gCNV tool (v4.3; gcnvkernel v0.8, ^19^), with an in-house background model for PCR-free WGS data.

**Figure 1.**
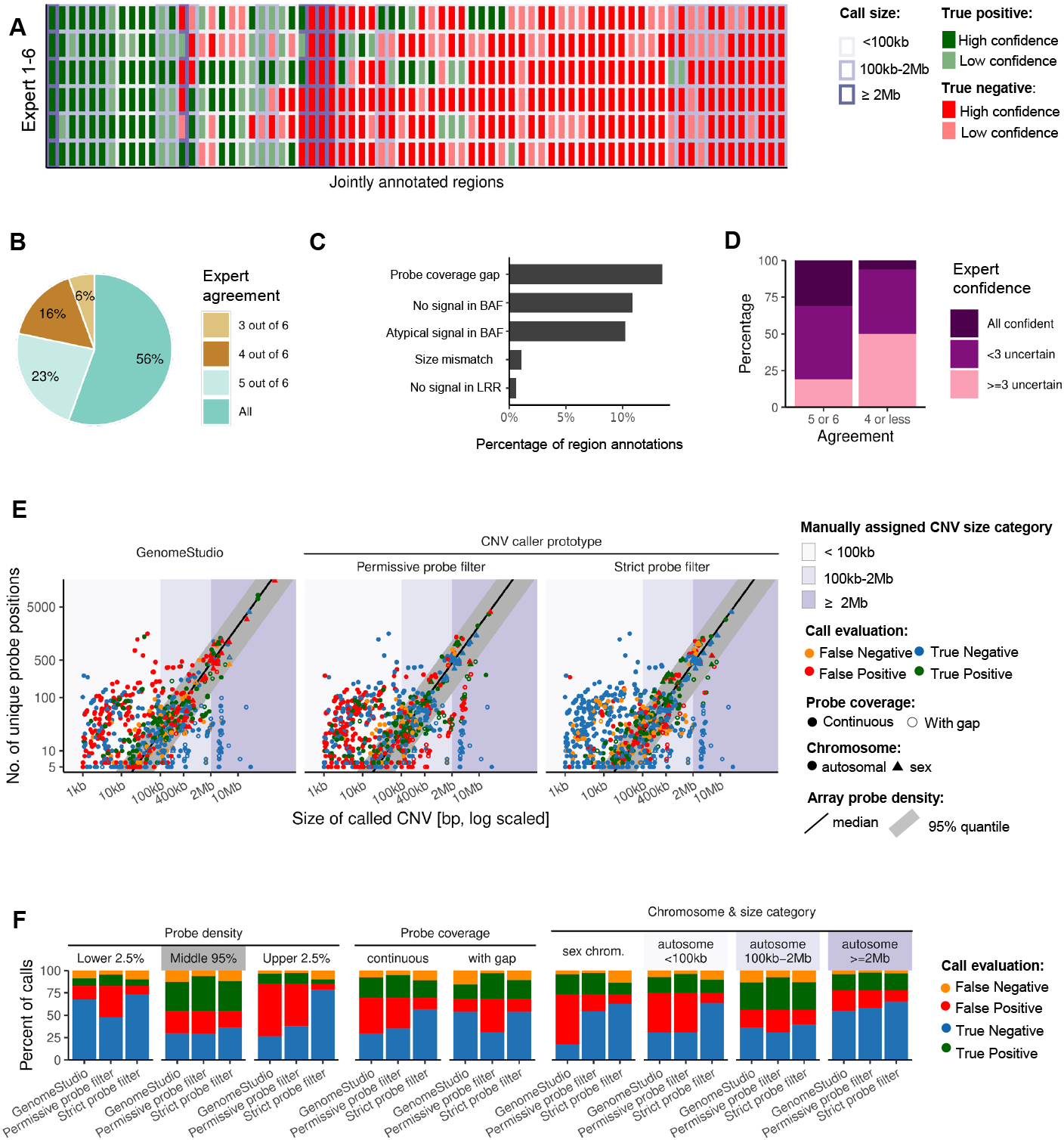
Expert evaluation identifies key strategies for enhancing CNV calling and detection. **(A)** Heatmap illustrating expert classifications of 74 jointly annotated CNV regions, with colours indicating true positive/negative calls and shading denoting confidence levels. X-axis: regions, ordered by hierarchical clustering of numerically scored evaluations (size bracket (1-3), scaled by confidence (0.5, 1) and negated for positive designations), Y-axis: individual experts. **(B)** Pie chart depicting the proportion of regions with expert agreement on the shared set in (A). **(C)** Horizontal bar chart summarizing the percentage of annotated regions with interpretive challenges across the complete data set as described in Y-axis. **(D)** Stacked bar plot representing the distribution of expert confidence levels (full, partial, low) across different agreement categories in shared data set. **(E)** Scatterplots visualizing the relationship between CNV size (log scale) and number of unique probe positions for *GenomeStudio* and *StemCNV-check* prototype under permissive and strict probe filter settings. **(F)** Grouped bar charts comparing the proportion of call classifications across probe density strata, probe coverage types, chromosome categories, and manually assigned size categories for each analysis workflow; quantifying data shown in (E).

**Figure 2.**
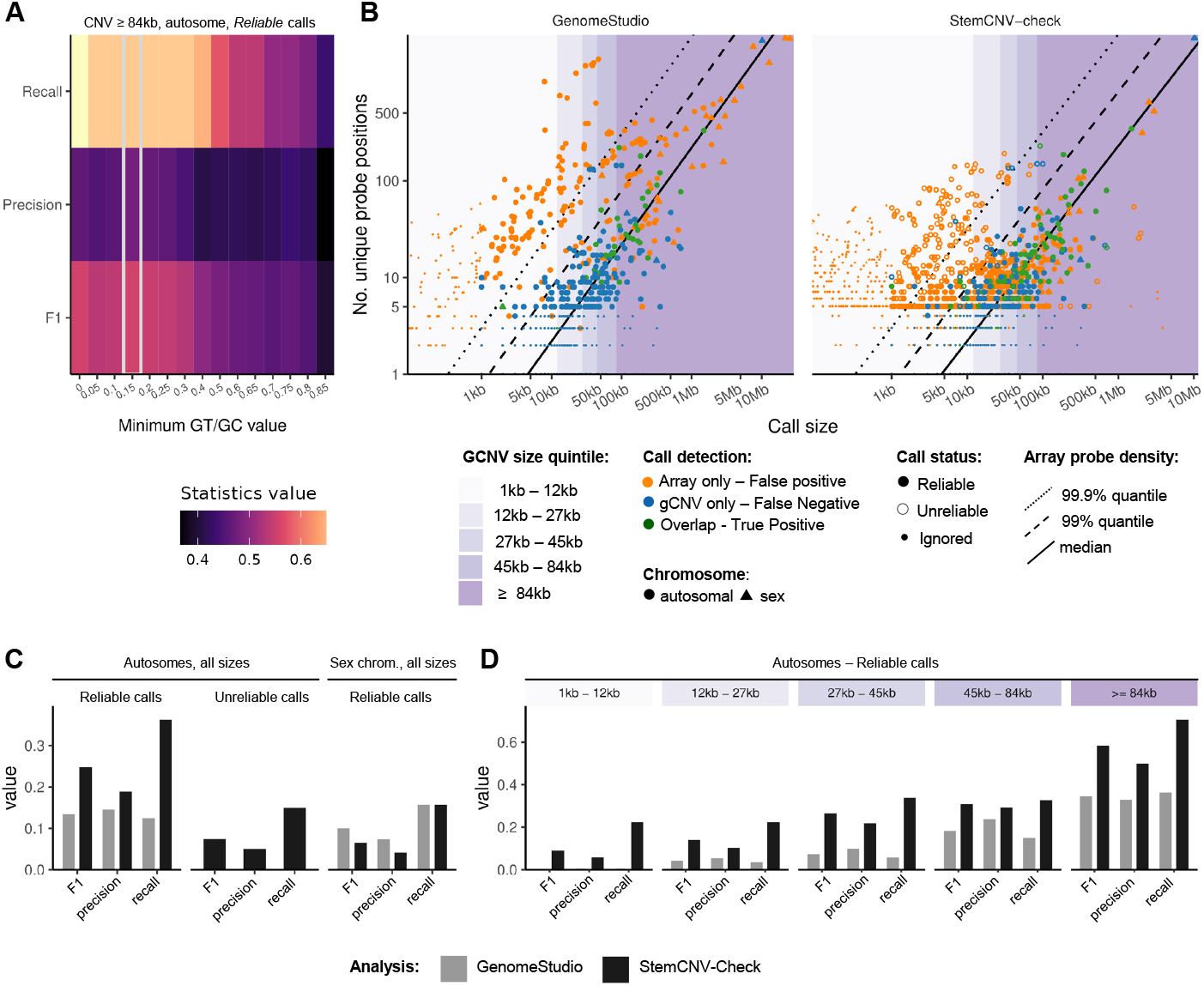
Benchmarking against WGS-derived CNVs shows that *StemCNV-check* outperforms *GenomeStudio*. **(A)** Heatmap depicting recall, precision, and F1 scores for *reliable* autosomal CNVs ≥84 kb across a range of GT/GC probe filter thresholds. The GT cut-off equals the GC cut-off. Grey box highlights the highest F1 score at GT/GC cut-off of 0.15. **(B)** Scatterplots illustrating CNV size versus number of unique probe positions for *GenomeStudio* and *StemCNV-check*, with points classified by detection type and shaped by chromosome category; shaded regions indicate probe density quantiles, and vertical lines mark gCNV size quintile boundaries. **(C)** Bar graphs comparing F1 score, precision, and recall of *GenomeStudio* and *StemCNV-check* for autosomal and sex chromosome CNVs. **(D)** Size-stratified bar graphs highlighting differences in F1 score, precision, and recall for reliable autosomal CNVs, emphasizing *StemCNV-check* improvements in the smallest and largest event categories. *F1=harmonic mean of precision and recall*.

### Evaluation of CNVs calls by experts

To investigate and refine CNV calling performance, hiPSC array samples (BIH Charité dataset) were analyzed using *GenomeStudio* with the cnvPartition plugin (**Supplementary Table 3**) and a prototype of *StemCNV-check* incorporating PennCNV and CBS algorithms. For *StemCNV-check*, two probe filtering modes were tested: *Permissive* and *Strict* (**Supplementary Table 3**). A union of all CNV calls across CNV callers and setting combinations (tools) was generated from all samples excluding those with call rates (percent of successfully genotype probes) below 0.99. Calls covering fewer than five distinct probes or less than 1 kb were removed. Overlapping calls were merged into non-overlapping regions and categorized by probe count (5–9, 10–49, ≥50) and size (1–10 kb, 10–100 kb, 100–400 kb, 400 kb–2 Mb, >2 Mb). Regions were also classified by workflow origin: only *GenomeStudio*, not *GenomeStudio*, or mixed. This resulted in 3 (probe count) ×5 (size) ×3 (workflow) region categories. Up to 13 gain and deletion examples were sampled per category, yielding 855 regions for manual review. Additional autosomal CNVs >2 Mb were included, as well as sex chromosome CNVs >2 Mb detected by at least 2 tools (e.g. PennCNV and CBS) to reduce spurious calls. In total, 928 regions were distributed to experts for evaluation.

A set of 74 shared regions, along with approximately 130 additional unique regions per expert, was manually evaluated by six experts. Each region was presented with calls from all individual tools or workflows, anonymized and randomly shuffled to reduce potential bias. Experts classified each call as a true CNV or a false positive. An initial unsupervised assessment was conducted to identify recurring error patterns and sources of ambiguity. Based on these insights, the final evaluation included detailed annotations of the following features: true positive call (yes/no), judgment uncertainty (N/A, too few probes, BAF unclear, other), signal mismatch (N/A, BAF no signal, LRR no signal), atypical BAF pattern (yes/no), presence of a probe gap (yes/no), and whether the true CNV extended beyond the called region (size mismatch yes/no) (**Supplementary Figure 1**). Experts also recorded clear tool-specific errors, such as size mismatches between the call and the presumed true CNV.

### Comparison of WGS and array analysis of matched samples

For benchmarking, we compared *StemCNV-check* v1.0 and Illumina *GenomeStudio* against CNV calls derived from WGS with matched SNP array from 19 samples: 13 in-house and 6 publicly available (Genome in a Bottle consortium -GIAB-https://www.nist.gov/programs-projects/genome-bottle) (**Supplementary Table 1 and 2**). CNV outputs from PennCNV, CBS, and integrated *StemCNV-check* results were evaluated under various probe filtering conditions. *GenomeStudio* (TSV) and gCNV (VCF) outputs were parsed for comparison. Probe filters were used with varying thresholds for the Illumina genotyping quality scores GenCall (GC) and GenTrain (GT) scores (v1-v15, **Supplementary Table 3**), with duplicate probe positions reduced to the highest GC score and pseudoautosomal and X-translocated (XTR) probes excluded from male samples.

CNVs from gCNV were annotated with the number of unique array probe positions overlapping each call, and with flag status according to criteria to enable evaluation of flag performance (**Table 1**). *GenomeStudio* calls were annotated with probe counts and minimum flags (to identify *ignored* call type). For array-based CNV evaluation, comparisons were made to WGS-derived gCNV calls. For each gCNV call, the percentage of overlap with one or more array calls was calculated, along with the reciprocal overlap of the array call(s) with the gCNV region. A reciprocal overlap exceeding 50% was considered sufficient to indicate replication of the gCNV call. Call size categories were determined using 20% quantiles of the gCNV call size distribution.

**Table 1.** CNV flag definitions and call reliability criteria. List of automated flags and resulting call types used in *StemCNV-check* to assess CNV quality and reliability.

| Flag | Description | Call type |  |  |
| --- | --- | --- | --- | --- |
|  |  | *Ignored* | *Unreliable* | *Reliable* |
| **min_size** (minimal size of CNV) | <1000bp | at least one | no | no |
| **min_probes** (minimal number of probes ) | <5 probes |  | no | no |
| **min_density** (minimal probe density per call size) | <10 per 1Mb |  | no | no |
| **high_probe_dens** (too high probe density per call size) | probe density >99% than the array | any | at least one | no |
| **probe_gap** (presence of a gap between probes) | no probe region between two probes<br>(min. 33% of call, more with low probe count) |  |  | no |

For calculation of recall, precision, and F1 score (harmonic mean of recall and precision) statistics, the data were stratified by multiple features. For each analysis workflow with its associated probe filter settings (*GenomeStudio* or *StemCNV-check* with various filters), *Reliable* autosomal CNV calls (without any flags) were split by original CNV caller (PennCNV, CBS, “combined-call” = intersection of both, or “*StemCNV-check*” = union of both), and size category. CNVs from sex chromosomes and autosomal *non-Reliable* CNVs were split by flag status (*Reliable/Unreliable/Ignored*) and original CNV caller, but not by size brackets to ensure sufficient counts per group for reliable summary statistics.

Comparative analyses were conducted using R v4.1.3, utilizing the *tidyverse* and *plyranges* packages.

### Design of the StemCNV-check workflow

*StemCNV-check* is implemented as a modular workflow using Snakemake (v8) ^20^ accompanied by a Python-based user interface that supports streamlined project setup and data preparation (publicly available on GitHub, https://github.com/bihealth/StemCNV-check). The workflow comprises several coordinated steps, described below.

#### Raw data processing

The pipeline begins with raw data processing using *iaap GenCall* (v1.1), followed by conversion of the output to VCF format via *gtc2vcf* (v1.16; https://github.com/freeseek/gtc2vcf).

Probe filtering is a key preprocessing step, as CNV callers are sensitive to noise from low-quality or redundant probes. While traditional methods use a simple GC score threshold, *StemCNV-check* supports more flexible filtering using both GC and GT scores. Users can customize filtering to: (1) manage duplicate probe positions, (2) handle pseudo-autosomal regions, and set minimum GC (3) and GT (4) thresholds— enabling cleaner input for downstream CNV detection (**Supplementary Table 3**).

Filter settings in *StemCNV-check* are defined via a configuration file. Based on benchmark analyses (**Figure 2A**), the recommended defaults (**Standard probe filter; Supplementary Table 3**) include: minimum GC and GT scores of 0.15, retaining only the highest GC-scoring probe at duplicate positions, and removing probes in pseudo-autosomal (PAR1, PAR2) and X-translocated (XTR) regions in male samples. Probes on the Y chromosome are always excluded for female samples. Genomic coordinates for these regions were sourced from Ensembl, with the XTR region defined by markers DXS1217–DXS3 on X and SY20/DXYS42– DXYS1 on Y ^21–23^. All filters are applied as soft-filters, so that data is masked but not lost. Following filtering, SNP genotypes in the VCF are annotated using *mehari* (v0.35; transcript database v0.10.3; https://github.com/varfish-org/mehari), reporting only the most severe consequence per allele, with prioritization of MANE-Select transcripts for consequence assignment. The final VCF file with probe data extends the initial *gtc2vcf* output with the probe-filters and mehari annotations.

#### CNV calling and processing

*StemCNV-check* employs both PennCNV (v1.0.5, ^24,25^) and circular binary segmentation (CBS) for CNV detection. PennCNV uses a Hidden Markov Model to infer copy number states from log R ratio (LRR) and B allele frequency (BAF), with default parameters except for enabling LOH detection. Analyses are run separately for autosomes, X, and Y chromosomes. CBS is implemented via the *DNAcopy* R/Bioconductor package (v1.76.0, ^26^) and a custom wrapper applies segmentation to LRR values. Segments exceeding predefined LRR thresholds are reported as CNVs. Adjacent CNV calls from each algorithm are merged using the default criteria: calls separated by <500 bp or <10 filtered probes, or calls overlapping when each is extended by 60% from its center.

Post-merging, CNVs failing quality thresholds are flagged (but retained) in the VCF if: <1,000 bp (“min_size”), supported by <5 probes (“min_probes”), or with probe density <10 probes/Mb (“min_density”) (**Table 1**). All flags are implemented as soft-filters within the VCF output. To generate a unified CNV call set per sample, overlapping calls from different algorithms are merged if at least one call covers ≥50% of the combined region and the minimum coverage by any single algorithm is ≥60%. Following integration, merged calls are re-evaluated under the same flag criteria. Each CNV is then annotated with additional features: (1) overlap with gene bodies (by default from GENCODE, release 45); (2) reciprocal overlap (≥50%) with CNVs from a defined reference sample, annotated as “matching_reference”; and (3) further flags are checked and applied based on array probe contents, specifically the presence of array-specific regions with high probe density (“high_probe_dens”) or known gaps in probe coverage (“probe_gap”) (**Table 1**).

#### CNV scoring and evaluation

In addition to standard CNV annotations, *StemCNV-check* computes a “*Check-Score*”—a composite metric integrating CNV size, copy number state, and overlap with gene and region annotations relevant to hPSC, cancer and dosage sensitivity ^27,28^. The base score is size- and CN-dependent, calculated as:

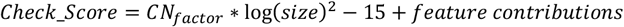

where *CN_factor* is 0.333 for CN = 1 or 3 (single copy gain/loss), 0.5 for CN = 0 or 4 (double loss/gain), and 0.275 for CN = 2 (copy-neutral LOH). Sex chromosome CN values are adjusted to reflect correct ploidy in male samples.

Additive feature contributions (values in brackets) are then included based on overlaps with annotated regions (**Figure 3B**): 1) CNV hPSC hotspots (**Supplementary Table 4**): genes (+30), sub-bands (+10), whole bands (+3); 2) dosage sensitive genes including haploinsufficiency and triplosensitivity (+5); 3) cancer driver genes (+5); 4) other genes (+0.2).

**Figure 3.**
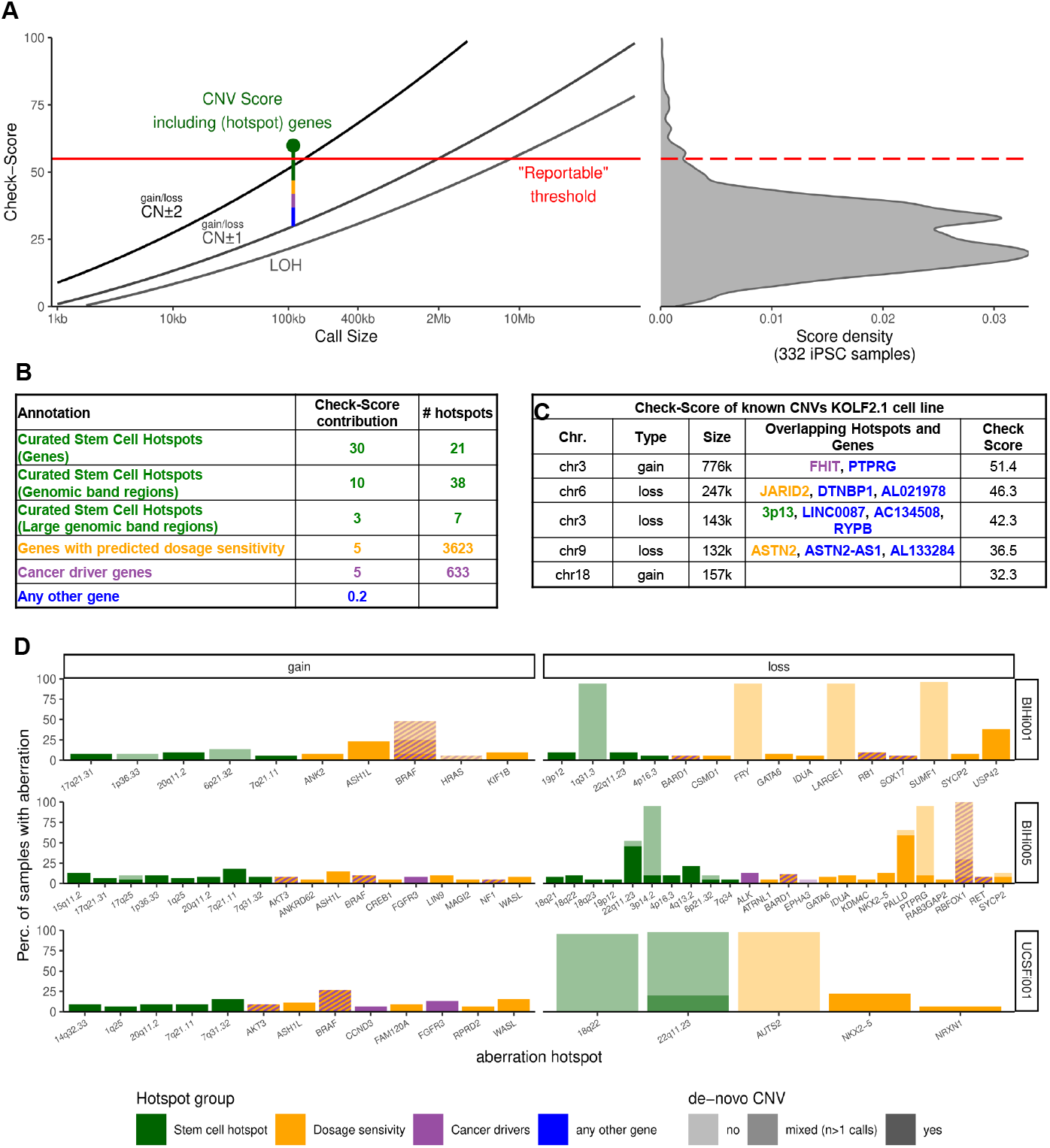
*Check-Score* framework for ranking CNVs by biological relevance. *(Left)* Schematic diagram illustrating the *Check-Score* calculation. Grey lines represent the base score as a function of CNV size and copy number (CN) deviation from neutrality, while the multi-colored bar depicts additional score contributions from overlapping annotation features. The red horizontal line marks the predefined threshold above which CNVs are classified as “*reportable*” or “*critical*”. *(Right)* Histogram presenting the density distribution of *Check-Scores* from 332 in-house hiPSC samples, including only *Reliable* CNVs derived using the standard probe filter. **(B)** Table summarizing the types of annotation features used in *Check-Score* calculation, their individual contribution values, and the number of features in each category. With the exception of general gene annotations, all features are CNV-type specific (gain/loss). **(C)** Table listing publicly reported CNVs from the KOLF2.1J cell line, showing overlaps with annotation features and the *Check-Score* values assigned by *StemCNV-check* for a representative sample. **(D)** Stacked bar plots visualizing the percentage of CNVs (gains or losses) overlapping *Check-Score* annotation features across three reference hiPSC line families. Only features with CNVs detected in at least three samples are displayed. Bars are segmented according to classification as *de novo*, reference-matching, or mixed (where part of the CNV overlaps with the reference and part is *de novo*, often in long genomic regions).

Each gene is scored once with its maximum applicable value. Scores are applied only when the CNV type matches the expected impact (gain or loss). CNVs overlapping user-defined regions of interest receive an additional 50-point score. Finally, calls are annotated with an estimated precision, derived from performance evaluations summarized in **Supplementary Table 5**.

Each CNV call is assigned a classification label based on its *Check-Score*, by call status (*Reliable/Unreliable/Ignored*) and reciprocal overlap (≥50%) with an optional reference sample (**Table 2A**). This labeling system prioritizes biologically relevant and high-confidence calls for interpretation while filtering out likely artifacts or background variation.

**Table 2.** CNV and SNV label definitions. **(A)** CNV label definitions and associated biological relevance. Classification scheme for CNVs in *StemCNV-check* reports, including label names, minimum *Check-Score* thresholds, match status to reference samples, and associated call type. Each label is linked to an interpretation of concern level. **(B)** SNV label definitions and criteria for classification. List of SNV categories used in *StemCNV-check* reports, specifying whether the variant matches the reference sample, relevant annotation categories (e.g., hotspot gene match, protein-altering effect), predicted impact (HIGH, MODERATE, or both), and the corresponding interpretation level of concern.

A. CNV labels
| CNV Label | Minimum *Check-Score* | Match in Reference Sample | Call Type | Concern / possible biological impact | Description |
| --- | --- | --- | --- | --- | --- |
| Critical de-novo | ≥ 55 | No | Reliable | High | High-confidence CNV call with potential biological relevance |
| Reportable de-novo | ≥ 55 | No | Unreliable | Medium | Reduced-confidence CNV call with potential biological relevance |
| De-novo call | < 55 | No | Reliable / Unreliable | Low | CNV call with unknown biological relevance |
| Reference genotype | any | Yes | any | None | CNV call that matches the reference sample genotype |
| Excluded call | any | No | Ignored | None | CNV call that does not match minimal quality requirements (close to "noise") |

| SNV Label | Match in Reference Sample | Categories (one or many) | Impact (Mehari annotation) | Interpretation/ Level of Concern | Description |
| --- | --- | --- | --- | --- | --- |
| **de-Novo Critical** | No | hotspot-match | MODERATE | High | SNV matches a described stem cell hotspot mutation, potentially affecting protein function. |
| **de-Novo Reportable** | No | hotspot-gene and/or | HIGH | Medium | SNV likely causes protein ablation in any gene (incl. gene with known stem cell hotspot mutation). |
|  |  | protein-ablation | MODERATE | Medium | SNV likely causes protein change in a gene with known stem cell hotspot mutation. |
| **de-Novo Unreliable critical/reportable** | No | hotspot-match /<br>hotspot-gene and/or<br>protein-ablation | HIGH / MODERATE | Low | Matches criteria for Critical or Reportable SNV label, but technical scores for genotype determination are low. If of concern, genotype should be confirmed by another method. |
| **de-Novo SNV** | No | other | HIGH / MODERATE | Low | SNV not present in reference genotype. Cause miss-sense mutations with potential impact but unknown effect on protein function. |
| **Reference genotype** | Yes | any | HIGH / MODERATE | None | Same genotype as detected in the reference sample. |

#### SNP & SNV analysis

*StemCNV-check* performs an identity check of the analyzed sample by comparing each sample to its reference (if defined), other samples on the same array chip, and those in the same sample group. Furthermore, additional samples to compare to can be defined via the config file. Genotype similarity is assessed using Manhattan distance across SNPs with high-confidence calls in all samples after filtering. Pairwise distances are calculated to identify divergence between each sample and its reference.

To identify SNPs likely to affect gene function, *StemCNV-check* uses *mehari* for transcript annotations. SNVs are listed showing heterozygous or homozygous genotype change compared to reference. Determined SNVs are further categorized based on their overlap with: (1) known recurrent mutations in hPSCs (2) genes frequently mutated in hPSCs, (3) HIGH-impact variants likely to disrupt protein function, or (4) variants with uncertain but potential impact, such as missense mutations or in-frame insertions/deletions (for (1) and (2) **Supplementary Table 4**). SNVs are then labeled to support interpretation: “reference genotype” if matching a reference sample; “de-novo critical” if falling into category 1; “de-novo reportable” if in category 2 or 3; “de-novo unreliable critical” if in categories 1–3 but with a GC score below 0.2 in the sample or reference; and “de-novo SNV” if none of the above apply (**Table 2B**).

### Report generation and additional outputs

*StemCNV-check* compiles all analysis results into an automatically generated, user-friendly HTML report using a built-in R Markdown template. In addition, it produces an Excel summary table containing basic information for all samples in the workflow run. As an alternative final output, users can generate a combined CNV table across all samples via command-line options. By default, this table includes all CNVs not labeled as “excluded call”.

## Results

### Expert evaluation identifies key strategies for enhancing CNV calling and detection

#### Expert Annotation and Dataset Creation

To overcome the limitations of *GenomeStudio*, restricted to cnvPartition algorithm and lacking flexible annotation or automation, we developed *StemCNV-check*, a modular pipeline integrating PennCNV and CBS for improved precision and recall (*StemCNV-Check* Workflow, **Figure 4**).

**Figure 4.**
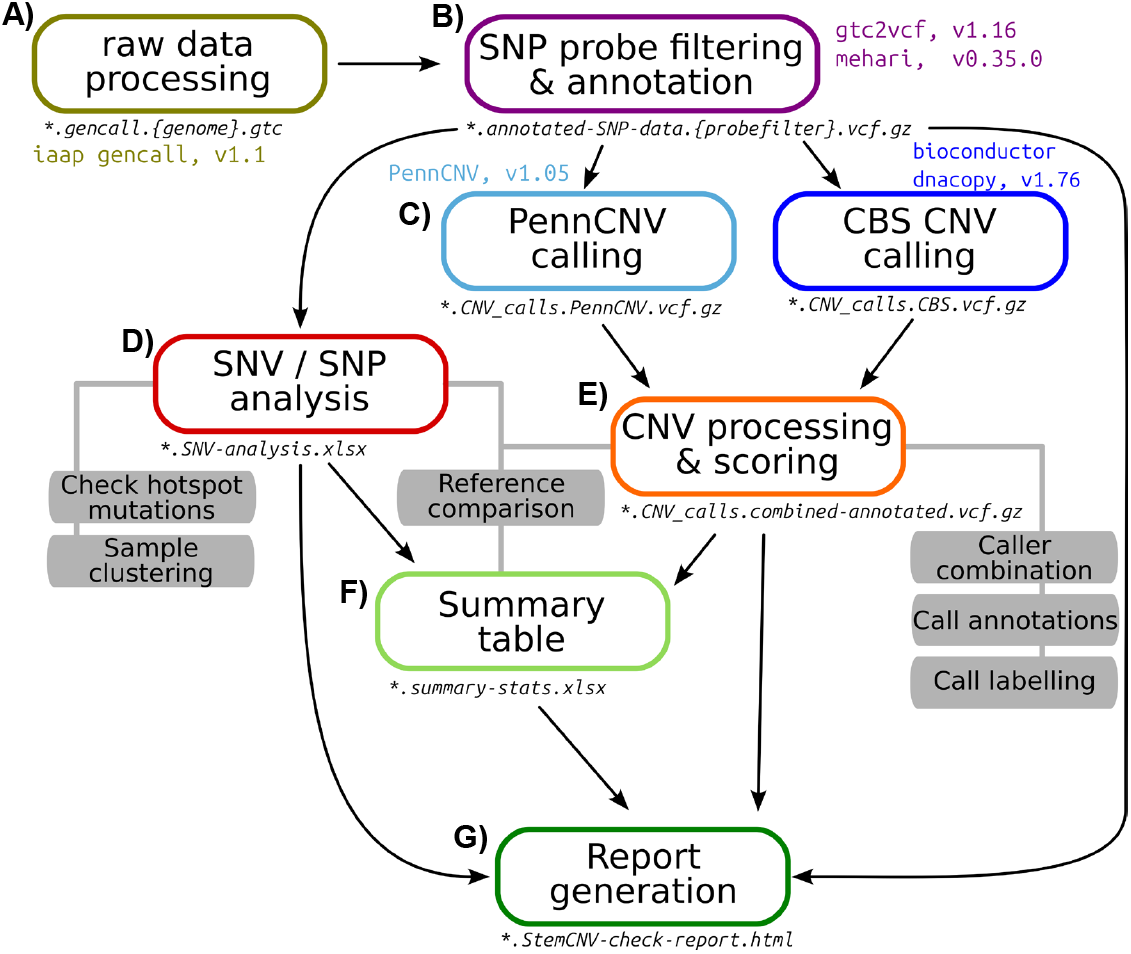
Workflow of the *StemCNV-check* pipeline. Graphic representation of the *StemCNV-check* workflow for a single sample. Colored boxes indicate steps generating permanent output files, with file patterns listed below; intermediate steps are omitted for clarity. Arrows show dependencies between processes. External tools or libraries are labeled with version numbers, while steps without version info correspond to custom R scripts. Grey boxes highlight key analyses at each stage. The workflow proceeds from **(A)** to **(G)** stepwise.

During early development, the *StemCNV-check* prototype caller generated many suspected false positives. To refine performance, we assembled a manually curated dataset of 928 candidate CNVs regions derived from both *GenomeStudio* and *StemCNV-check* prototype, the latter applied with varying probe filter settings (see Methods, **Supplementary Table 3**). Six independent experts evaluated these regions using BAF and LRR plots comparing tools and filtering strategies under blinded conditions (see Methods, **Supplementary Figure 1**).

The dataset was designed for tool optimization rather than representing the genomic CNV landscape. Regions were deliberately selected to span a broad and balanced range of CNV sizes, probe densities and signal complexities, enabling systematic evaluation of CNV calling behavior across diverse scenarios.

To assess inter-expert agreement and establish a reference baseline, a subset of 74 regions was jointly reviewed and annotated (**Figure 1A**). Among these shared regions, 56% reached full consensus, 23% showed one dissenting expert, and 22% had two or more divergent classifications (**Figure 1B**). Consensus varied with CNV size. Agreement was higher for CNVs >100 kb but declined for smaller events (≤100 kb), often due to weak signals or sparse probe coverage.

Experts were also asked to annotate qualitative features that could influence interpretability. Qualitative annotations highlighted challenges: 13.5% of regions had probe coverage gaps, 10.9% no signal in BAF, and 10.2% atypical BAF signals (**Figure 1C**). When experts were unable to confidently classify a region, due to noise, insufficient resolution, or conflicting evidence, this uncertainty was explicitly recorded.

Confidence in CNV interpretation correlated with expert agreement: regions with broad consensus were classified with high or good confidence in ∼80% of cases, whereas discordant regions showed low confidence in half of evaluations (**Figure 1D**).

#### Analysis of expert annotations

Given the strong expert consensus (**Figure 1D**) and consistent confidence patterns between the shared and full sets (**Supplementary Figure 2A**), we used the full dataset as ‘ground truth’. This enabled comparison of *GenomeStudio* (cnvPartition) with the *StemCNV-check* prototype under two probe filtering modes: *Permissive* (Illumina’s minimum recommendation for reliable genotypes ^29^ and *Strict* (retaining only higher-quality probes; **Supplementary Table 3**). Both filters ensured one probe per genomic position but differed in quality thresholds. Across filters, *StemCNV-check* prototype consistently avoided false positives in high-density regions frequently misclassified by *GenomeStudio* (**Figure 1E**). CNVs within the central 95% probe density were reliable across methods, whereas gaps or low probe density increased false positives, especially with the *Permissive* filter (**Figure 1F**). In regions with continuous coverage, *GenomeStudio* produced more false positives than *StemCNV-check* prototype; in gap-containing regions, *Strict* probe filtering and *GenomeStudio* performed similarly, while *Permissive* probe filtering yielded more false positives (**Figure 1E+F**). Performance was size-dependent: >100 kb calls were similar across tools, but the *Strict* filter improved reliability for smaller events (<100 kb) (**Figure 1F**). Overall, *StemCNV-check* with *Strict* filtering showed higher specificity and reliability across sizes and probe densities, outperforming *GenomeStudio* in difficult regions.

*StemCNV-check* applies a tiered classification to improve consistency beyond probe filtering. Initially, three features (small size, low probe count, low probe density) were used to flag technically uninterpretable calls, marked as *Ignored* but retained for transparency. Expert evaluation revealed additional recurrent error sources: probe gaps and regions of unusually high density, leading to the introduction of *Unreliable* flags. Calls passing all criteria were labeled *Reliable*. This system standardizes call interpretation while preserving problematic calls for review (see Methods and **Table 1**).

### StemCNV-check outperforms GenomeStudio in precision with comparable recall

We next benchmarked CNV calls against matched WGS in 19 samples (**Supplementary Table 1-2**). To identify *StemCNV-check* settings for optimal performance, we evaluated recall, precision, and the F1 score across a range of probe filter thresholds. Optimal probe filters (termed “*Standard”*) were chosen based on the highest F1 score for *Reliable* autosomal calls falling within the largest 20% of WGS-derived events (this are the CNVs ≥84 kb) (**Figure 2A, Supplementary Table 3**). Comparison with gCNV confirmed that high-density CNVs are largely false positives (**Figure 2B**), supporting the need to separate *Reliable* from *Unreliable* calls in *StemCNV-check*.

Reliable autosomal CNVs from *StemCNV-check* outperformed *GenomeStudio* across recall, precision, and F1, while sex chromosome performance was comparable. Unreliable CNVs had recall similar to *GenomeStudio* but lower precision and F1, indicating more false positives (**Figure 2C**).

Stratification of reliable calls by size showed *StemCNV-check* outperforming *GenomeStudio* for all sizes with the largest improvements for small (≤45 kb) and large (≥84 kb) CNVs (**Figure 2D**). *GenomeStudio* generated more false positives, including >400 kb events on sex chromosomes which were associated with average probe density (**Figure 2B, Supplementary Figure 3A**).

To further assess performance, we stratified CNVs by their caller origin (PennCNV only, CBS only, combined, or the full *StemCNV-check* call set). CNVs detected by both PennCNV and CBS consistently showed the highest precision, reflecting their greater likelihood of being true positives (**Supplementary Figure 3B, Supplementary Table 5**). The precision values from this performance analysis are now also incorporated into *StemCNV-check* as annotations. This addition provides users with a probabilistic reliability estimation for each CNV, extending beyond binary flagging and offering a quantitative measure of confidence. Integrating caller overlap with precision estimates illustrates how *StemCNV-check’s* multilayered approach improves CNV accuracy, especially in autosomal regions.

#### Sex chromosome call handling

CNV detection on sex chromosomes presents specific challenges due to hemizygous content and atypical probe behavior. In our evaluation, *StemCNV-check* consistently showed higher reliability than *GenomeStudio* across filter settings, particularly with the *Strict* probe filter (**Supplementary Figure 2B**), indicating better noise handling and signal discrimination. However, performance gains were not uniform across CNV sizes, and in some cases, accuracy was comparable between tools (**Figure 2C**). Interpretation was further limited by the small number of gCNV calls on sex chromosomes, preventing robust size-stratified benchmarking. In addition, *GenomeStudio* and *StemCNV-check* differ in their handling of large sex chromosome CNVs and *StemCNV-check’s Unreliable* category captures many ambiguous calls (**Supplementary Figure 3A**). Together, these findings show that targeted filtering improves detection on sex chromosomes but also underscore the need for further optimization in these regions.

### Check-Score ranks CNVs based on their biological relevance and level of concern

#### Check-Score development

While *StemCNV-check* sensitively and robustly detects CNVs, size alone is insufficient to judge biological impact, as gene- and region-specific effects strongly influence hPSC function and stability.

To address this, we developed *Check-Score*, integrating CNV size with genomic and functional annotations to prioritize biologically meaningful events.

The framework was informed by a literature review of 40 studies to compile recurrent hPSC CNVs (**Supplementary Table 4**, https://bihealth.github.io/StemCNV-check/CNV-hotspots/index_1.html). These included regions recurrently gained or lost and genes linked to hPSC growth or fate. To extend beyond stem cell hotspots, we added cancer driver genes (IntOGen) ^28^ and genome-wide dosage sensitivity predictions^27^, creating a comprehensive feature set.

*Check-Score* begins with a size- and copy-state–adjusted base value, then adds weighted contributions for biologically relevant features: stem cell hotspots, dosage-sensitive genes, cancer drivers, and other genes (**Figure 3A-B**). Hotspot, dosage, and cancer annotations are CNV-type-specific (gain or loss), whereas general gene annotations apply regardless of CNV type.

We applied *Check-Score* to categorize CNVs and designed a labeling system for systematic interpretation (**Figure 3A, Table 2A**). Ideally, each hPSC sample is compared to a matched reference (e.g., donor fibroblasts or earliest passage), so that CNVs absent in the reference are classified as *de novo*. If no reference is available, all CNVs default to *de novo* label. Detected CNVs are ranked by *Check-Score*, integrated with reference status and *StemCNV-check* quality flags. Events exceeding the predefined threshold of **55**, derived from prior QC experience to ensure CNVs >2 Mb are flagged, are highlighted as *reportable* or *critical*. The distinction reflects reliability: critical calls pass all quality criteria, while reportable calls carrying flags, such as probe gaps or high probe density, are *Unreliable* indicating possible technical artifacts. This framework prioritizes high-confidence, biologically relevant CNVs while retaining transparency on calls requiring caution.

#### Check-Score validation

Validation with the KOLF2.1J hiPSC line, described as exhibiting high genomic stability, robust differentiation capacity, and resilience to genetic engineering, and therefore proposed as a reference standard for hiPSC research^30^, confirmed known CNVs on chromosomes 3, 6, 9, and 18 (**Figure 3C**). None exceeded the reportability threshold, consistent with prior studies^31,32^, but the framework highlighted overlaps with biologically relevant genes, including *FHIT* (cancer driver) and the predicted dosage-sensitive *JARID2* and *ASTN2*, illustrating its ability to annotate biologically relevant features.

Overall, *StemCNV-check* sensitively detects CNVs, benchmarks them against a reference, and ranks their biological relevance via *Check-Score*. Automated scoring offers a strong first assessment, but expert review remains essential. The tool is particularly valuable for monitoring recurrent alterations across passages and gene-editing events, providing critical insight into hPSC genomic stability.

#### Check-Score identifies recurring culture acquired CNVs

To assess *Check-Score* in practice, we analyzed 332 hiPSC lines, including gene-edited derivatives. Most CNVs scored below the reportability threshold, confirming a generally stable genomic landscape under standard culture (**Figure 3 A, D**). Notably, *Check-Score* highlighted recurrent *de novo* CNV gains at stem cell hotspots and cancer-associated loci across several line families. Within this cohort, we identified *de novo* CNV gains at known stem cell hotspot regions on chromosomes 20q11.2 and 7q21.11, as well as at the oncogene *BRAF*, across all three evaluated cell line families—BIHi001, BIHi005, and UCSFi001—with observed frequencies ranging from about 10% to 30% (**Figure 3D**). These CNV events were detected in gene-edited clones and in both, master and working banks of gene-edited cells, when compared to their earliest available parental controls. Additional CNV gains were observed at 17q21 in both the BIHi001 and BIHi005 families, while a gain at 7q31.32 was shared between BIHi005 and UCSFi001 (**Figure 3D**). A loss at 1q31.3 was identified in the reference sample from the BIHi001 family; however, this was not considered a *de novo* event in subsequent family-derived samples (**Figure 3D**). Notably, approximately 44% of samples from the BIHi005 family exhibited a *de novo* loss at 22q11.23. While CNVs were also detected in the same region within the UCSFi001 family, they were not classified as *de novo*. In a subset of samples, CNV detection in this region yielded multiple segmented calls. These were manually reviewed and validated by expert curators, indicating variability in call resolution and analysis quality (**Figure 3D**).

#### SNV/SNP analysis allows sample identification and highlights genotype changes with potential biological impact

*StemCNV-check* performs SNP analysis by comparing genotype similarity among samples in the same run. In a cohort reprogramming experiment (30 hiPSC clones from 6 donors), SNP distance matrices and hierarchical clustering correctly matched clones to their donors, revealing over-representation of one donor in the subset of samples analyzed and identified mixed clones (**Supplementary Figure 4**).

Additionally, *StemCNV-Check* lists coding SNVs differing from the reference genome as *de novo*, while shared variants are reported separately. Each SNV is labeled as critical, reportable, unreliable, or general *de novo* (**Table 2B**), based on hotspot overlap and predicted protein impact (*mehari* annotation; **Supplementary Table 4**). This labeling supports assessment of unwanted SNV risk.

### StemCNV-check Workflow

Building on these improvements, we finalized *StemCNV-check* as a streamlined workflow for assessing CNV impact in hPSCs (**Figure 4**). The pipeline starts with raw array processing and probe filtering/annotation (**Figure 4A-B**), followed by parallel CNV calling with PennCNV and CBS, integration, scoring, and reliability flagging (**Figure 4C**; **Table 1**). Calls are annotated for genomic context and merged by biological relevance (**Figure 4E**), while SNP analysis supports clustering and hotspot detection (**Figure 4D**). Outputs are consolidated into summary tables and an interactive HTML report (**Figure 4F-G**).

### StemCNV-check provides interactive reports for efficient and comprehensive CNV evaluation

*StemCNV-check* automatically generates an interactive HTML report that summarizes results and guides interpretation (**Figure 5**; **Supplementary HTML file**). The report includes **Sample overview, CNV calling, SNV analysis, and Sample comparison**, each with descriptive outputs to support user interpretation (**Figure 5, Supplementary Figure 5A**).

**Figure 5.**
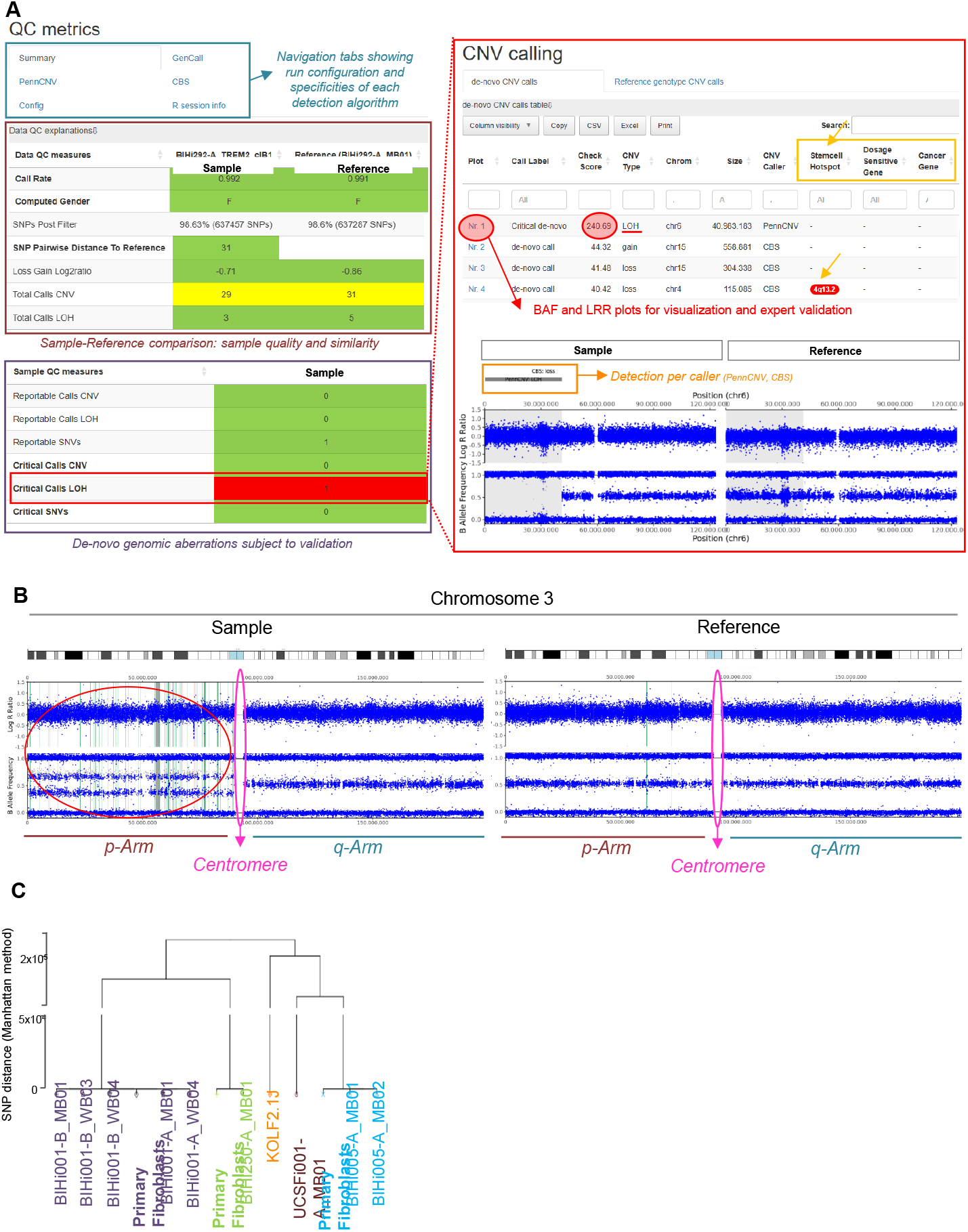
*StemCNV-check* provides interactive reports for efficient and comprehensive CNV evaluation. *(Left)* Interface screenshot of the quality control (QC) metric overview table with color-coded indicators (green, yellow, red) for sample quality, observed CNVs, and CNVs/SNVs classified as reportable or critical. *(Right)* Interactive CNV tables list either all *de novo* or reference-matching CNVs for a sample, including specifications and annotations, each entry links to a detailed call view. Associated CNV plots display LRR and BAF signals alongside genomic bands and gene tracks, with annotated hotspots highlighted. Separate tables list all overlapping genes and hotspots, with links to REEV, NCBI, Ensembl, and relevant publications. Depicted example: side-by-side LOH in chromosome 6 in a gene-edited hiPSC clone compared to its parental reference. Interface screenshot showing genome-wide and chromosome-specific plots for a large gain on chromosome 3; centromeric regions without probes are excluded from analysis. Red circle at p-arm indicates multiple call detection for a large CNV. **(C)** Interface screenshot of a dendrogram based on SNP distance, illustrating family-specific grouping (families indicated by colors) and detection of potential sample swaps.

**Sample overview** displays run details and side-by-side comparison with a given reference. QC metrics include SNP distance, gender match, and total calls, with *de novo* CNVs/SNVs flagged as reportable or critical (**Table 2A**). Visual alerts (e.g., red marking) highlight concerning findings (**Figure 5A**).

**CNV calling** ranks *de novo* CNVs by *Check-Score* in sortable tables with labels, type, size, precision, and probe features. Each entry links to side-by-side BAF/LRR plots, a unique and relevant feature for call validation by expert, and annotated gene lists with external resources (**Supplementary HTML file**). Parameters involved in *Check-Score* output (stem cell hotspot, dosage sensitive gene or cancer gene matching the CNV) are described so that potential biological impact of the CNV can be assessed. **Figure 5A** illustrates a large LOH on chromosome 6 in a CRISPR-edited clone, showing the value of post-editing QC. CNVs overlapping a region of interest (ROI) are also marked in the overview table. Reference matching CNVs and if defined regions of interest (ROI) are displayed in separate tabs.

**SNV analysis** lists coding variants relative to the reference, sorted by SNV label (**Table 2B**). Each row includes genotype change, ROI overlaps, gene affected and predicted protein effect (*mehari* annotation; **Supplementary Figure 5A**). Reference matching SNVs are displayed in separate tabs.

**Sample comparison** provides a genome-wide overview with side-by-side plots of sample and reference, aiding validation of large events (e.g., chromosome 3 gain detected as multiple smaller calls; **Figure 5B**). SNP-based clustering generates dendrograms that group related lines with possibility to spot sample swaps (**Figure 5C**). This feature is a fast, efficient and low-cost method to identify samples and possible swaps during culture handling, therefore, a suitable alternative to short tandem repeat (STR) based fingerprinting methods. We have determined threshold SNP distances based on the distribution observed in the cohort of samples analysed in this study. Thresholds defined from cohort data distinguish same-donor samples (≤100), LOH-associated changes (500–5000), and likely swaps (>5000; **Supplementary Figure 5B**).

Overall, the *StemCNV-Check* report delivers an intuitive, structured overview that streamlines CNV/SNV interpretation and enables routine monitoring of hPSC genomic stability, making it a powerful QC tool.

## Discussion

Karyotyping has long served as the standard for detecting large-scale genomic abnormalities but lacks the resolution to detect smaller, sub-chromosomal changes or SNVs. WGS offers strong sensitivity for all variants, yet cost, turnaround, and compute remain higher than with any other established method. While low coverage WGS approaches are being investigated, their analysis and interpretation are still difficult. Therefore, SNP arrays remain a very cost-effective choice for monitoring genomic stability via CNV and SNP analysis. *StemCNV-check* was designed for this fast-moving landscape: while optimized for SNP array data, its modular architecture and reliance on community standards allows easy extension to WGS based methods, while retaining other advancements. It delivers a full solution - detection, analysis, interpretation - that distinctively addresses the lack of hPSC specific tools, which so far impeded biological prioritization^33–36^. At the caller level, we implement a consensus strategy combining PennCNV and CBS, two complementary, actively maintained open-source algorithms, since such a combination can increase overall performance^37,38^. Together with advanced annotation and a biologically informed scoring system, *StemCNV-check* reduces false positives, preserves high sensitivity, and delivers an interactive report. Expert CNV annotation and evaluation highlighted the need to consider both probe input quality and CNV characteristics, leading to the development of a call flagging system that improves reliability while minimizing sensitivity loss. In its current version, *StemCNV-check* is optimized for large autosomal CNVs, achieving superior results compared with *GenomeStudio*. Benchmarking against WGS-derived CNVs confirmed this advantage, showing a better precision and recall, particularly in autosomal chromosomes. Further assessment was constrained by the limited number of true positive WGS reference calls, especially for sex chromosomes and flagged array regions, which prevented detailed evaluation across CNV sizes. Overcoming these limitations will require larger, well-annotated datasets and closer study of how array design influences CNV calling. Further improvements may also involve region-specific probe filters or parameter settings, technically challenging to implement and especially benchmark but likely more effective than a single global setting. Potential extensions could include incorporating CNV approaches from tumor genomics, which enable direct comparisons with clonally related normal samples^39^. Other high-performing tools, such as MoChA^40,41^, rely on cohort-based methods that present challenges for implementation and reproducibility when processing individual samples or batches in QC pipelines. In summary, several alternative CNV callers offer valuable opportunities to expand *StemCNV-check’s* capabilities, but their integration will require careful evaluation of methodological constraints, implementation complexity, and reproducibility in routine workflows.

*StemCNV-check* is a modular Snakemake workflow engineered for adaptability—scaling from workstations to clusters and able to readily accommodate new CNV callers, annotations, or future WGS modules. It is open source and unit tested, with a community-oriented design that encourages contributions and portability as we move forward. Furthermore, *StemCNV-check* is designed with accessibility in mind: users without bioinformatics expertise can readily interpret the comprehensive HTML report, and with minimal training, operate the workflow. Although the initial setup currently requires Linux support, we provide guidance for installation on modern Windows systems (https://stemcnv-check.readthedocs.io) and are working toward full platform independence.

A key innovation of *StemCNV-check* is the *Check-Score*, a biologically informed ranking system tailored to hPSCs. We acknowledge that the functional form and constants were empirically calibrated rather than derived from a theoretical model. This is a deliberate, transparent compromise: the score is not a directly interpretable value, but a reproducible approximation designed to prioritize biologically relevant CNVs. Importantly, feature annotations add explanatory value beyond size and copy number alone and provide a modular scaffold that can incorporate additional or alternative annotations without altering the core ranking principles. hPSC related specific features have been carefully curated in a database of recurrent genomic aberrations (hotspots) known to affect hPSC proliferation, pluripotency, or undifferentiated state, and is aligned with ISSCR guidelines for human stem cell research ^1^ and is made available as an online version (https://bihealth.github.io/StemCNV-check/CNV-hotspots/index_1.html) ^44^. Additional features including cancer-related genes and dosage-sensitive loci contribute to highlight variants with potential biological impact. This facilitates the identification of hPSC lines carrying detrimental changes, thereby strengthening QC and guiding sample selection for downstream applications.

Another *StemCNV-check* novelty lies in the SNP-based identity check, which enables easy detection of sample swaps and contamination, further assisting researchers in maintaining data integrity.

Beyond QC, *StemCNV-check* also has the potential to streamline analysis of hPSC data and enable the discovery of previously overlooked links between experimental conditions and CNV emergence, offering new insights into the mechanisms driving genomic instability in hPSCs.

Future incorporation of AI–based classification could further improve CNV prioritization, while adaptation for multi-omics data integration may extend its applicability beyond array-based technologies. Together, these directions position *StemCNV-check* as a new standard for SNP array–based CNV analysis, enabling more reliable, reproducible, and biologically informed stem cell research.

## Supporting information

Example report

## Availability

*StemCNV-check* is opensource (https://github.com/bihealth/StemCNV-check), easy to install via Bioconda (https://anaconda.org/bioconda/stemcnv-check), and fully documented with step-by-step guides for first time users (https://stemcnv-check.readthedocs.io). An example dataset is provided for quick testing and benchmarking. *StemCNV-check* is used in the authors’ daily work, actively maintained, and we welcome questions and suggestions via GitHub issues or email.

## Acknowledgements

We thank Dr. Jennifer Volz-Glazer, Institute of Molecular Biotechnology (IMBA), Vienna, Austria and Dr. rer. nat. Lukas Cyganek, Fraunhofer-Institut für Translationale Medizin und Pharmakologie (ITMP), Göttingen, Germany; for providing SNP array data of samples and complementary metadata. We thank the whole team of the Core Unit pluripotent Stem cells and Organoids (CUSCO) for supporting sample collection and analysis. We thank Carolin Genehr (MDC) who has generated data from cohort reprogramming experiment. We thank all early users of *StemCNV-check* for their valuable feedback and bug reports.

## Author contributions

**N.v.K**. designed methods, implemented algorithms, developed analysis tools, performed computational analyses, contributed to expert annotation, published resources on GitHub, and wrote the manuscript and user documentation. **V.F.V**. acquired and curated data, analyzed and validated results, contributed to expert annotation, curated the hotspot list, co-wrote the manuscript, and reviewed user documentation. **K.A.L**. acquired and curated data, analyzed and validated results, contributed to expert annotation, co-wrote the manuscript, and reviewed user documentation. **M.B**. performed data analysis and contributed to user documentation. **S.D**. tested the tool, assisted in data acquisition, contributed to expert annotation and reviewed the manuscript. **H.S**. and **D.B**. conceived the study, contributed to method design and supervision, secured funding, and reviewed the manuscript and user documentation. **H.S**. performed testing of tool prototypes and contributed to expert annotation. **All authors** reviewed and approved the final manuscript and documentation.

## Declaration of generative AI and AI-assisted technologies in the writing process

During the preparation of this work the authors used ChatGPT in order to shorten the draft and increase readability and language. After using this tool, the authors reviewed and edited the content as needed and take full responsibility for the content of the published article.

## References

1. Ludwig, T.E., Andrews, P.W., Barbaric, I., Benvenisty, N., Bhattacharyya, A., Crook, J.M., Daheron, L.M., Draper, J.S., Healy, L.E., Huch, M., et al. (2023). ISSCR standards for the use of human stem cells in basic research. Stem Cell Rep. 18, 1744–1752. 10.1016/j.stemcr.2023.08.003.

2. Andrews, P.W., Barbaric, I., Benvenisty, N., Draper, J.S., Ludwig, T., Merkle, F.T., Sato, Y., Spits, C., Stacey, G.N., Wang, H., et al. (2022). The consequences of recurrent genetic and epigenetic variants in human pluripotent stem cells. Cell Stem Cell 29, 1624–1636. 10.1016/j.stem.2022.11.006.

3. Assou, S., Girault, N., Plinet, M., Bouckenheimer, J., Sansac, C., Combe, M., Mianné, J., Bourguignon, C., Fieldes, M., Ahmed, E., et al. (2020). Recurrent Genetic Abnormalities in Human Pluripotent Stem Cells: Definition and Routine Detection in Culture Supernatant by Targeted Droplet Digital PCR. Stem Cell Rep. 14, 1–8. 10.1016/j.stemcr.2019.12.004.

4. Spits, C., Mateizel, I., Geens, M., Mertzanidou, A., Staessen, C., Vandeskelde, Y., Van der Elst, J., Liebaers, I., and Sermon, K. (2008). Recurrent chromosomal abnormalities in human embryonic stem cells. Nat. Biotechnol. 26, 1361–1363. 10.1038/nbt.1510.

5. Amps, K., Andrews, P.W., Anyfantis, G., Armstrong, L., Avery, S., Baharvand, H., Baker, J., Baker, D., Munoz, M.B., Beil, S., et al. (2011). Screening ethnically diverse human embryonic stem cells identifies a chromosome 20 minimal amplicon conferring growth advantage. Nat. Biotechnol. 29, 1132–1144. 10.1038/nbt.2051.

6. Laurent, L.C., Ulitsky, I., Slavin, I., Tran, H., Schork, A., Morey, R., Lynch, C., Harness, J.V., Lee, S., Barrero, M.J., et al. (2011). Dynamic Changes in the Copy Number of Pluripotency and Cell Proliferation Genes in Human ESCs and iPSCs during Reprogramming and Time in Culture. Cell Stem Cell 8, 106–118. 10.1016/j.stem.2010.12.003.

7. Al Delbany, D., Ghosh, M.S., Krivec, N., Huyghebaert, A., Regin, M., Duong, M.C., Lei, Y., Sermon, K., Olsen, C., and Spits, C. (2024). De Novo Cancer Mutations Frequently Associate with Recurrent Chromosomal Abnormalities during Long-Term Human Pluripotent Stem Cell Culture. Cells 13, 1395. 10.3390/cells13161395.

8. Avior, Y., Lezmi, E., Eggan, K., and Benvenisty, N. (2021). Cancer-Related Mutations Identified in Primed Human Pluripotent Stem Cells. Cell Stem Cell 28, 10–11. 10.1016/j.stem.2020.11.013.

9. Gore, A., Li, Z., Fung, H.-L., Young, J.E., Agarwal, S., Antosiewicz-Bourget, J., Canto, I., Giorgetti, A., Israel, M.A., Kiskinis, E., et al. (2011). Somatic coding mutations in human induced pluripotent stem cells. Nature 471, 63–67. 10.1038/nature09805.

10. Lei, Y., Al Delbany, D., Krivec, N., Regin, M., Couvreu De Deckersberg, E., Janssens, C., Ghosh, M., Sermon, K., and Spits, C. (2024). SALL3 mediates the loss of neuroectodermal differentiation potential in human embryonic stem cells with chromosome 18q loss. Stem Cell Rep. 19, 562–578. 10.1016/j.stemcr.2024.03.001.

11. Lezmi, E., and Benvenisty, N. (2021). Identification of cancer-related mutations in human pluripotent stem cells using RNA-seq analysis. Nat. Protoc. 16, 4522–4537. 10.1038/s41596-021-00591-5.

12. Merkle, F.T., Ghosh, S., Kamitaki, N., Mitchell, J., Avior, Y., Mello, C., Kashin, S., Mekhoubad, S., Ilic, D., Charlton, M., et al. (2017). Human pluripotent stem cells recurrently acquire and expand dominant negative P53 mutations. Nature 545, 229–233. 10.1038/nature22312.

13. Haake, J., and Steenpass, L. (2025). Chromosomal quality control in hPSCs: A practical guide to SNP array analysis with GenomeStudio. Front. Cell Dev. Biol. 13, 1599923. 10.3389/fcell.2025.1599923.

14. Lin, C.-F., Naj, A.C., and Wang, L.-S. (2013). Analyzing Copy Number Variation using SNP Array Data: Protocols for Calling CNV and Association Tests. Curr. Protoc. Hum. Genet. Editor. Board Jonathan Haines Al 79, Unit-1.27. 10.1002/0471142905.hg0127s79.

15. Montalbano, S., Sánchez, X.C., Vaez, M., Helenius, D., Werge, T., and Ingason, A. (2022). Accurate and Effective Detection of Recurrent Copy Number Variants in Large SNP Genotype Datasets. Curr. Protoc. 2, e621. 10.1002/cpz1.621.

16. Chue Hong, N.P., Katz, D.S., Barker, M., Lamprecht, A.-L., Martinez, C., Psomopoulos, F.E., Harrow, J., Castro, L.J., Gruenpeter, M., Martinez, P.A., et al. (2021). FAIR Principles for Research Software (FAIR4RS Principles). 10.15497/RDA00068.

17. Fischer, I., Küchler, J., Schaar, C., Fisch, T., Cernoch, J., Fischer, K., Fernández Vallone, V., Kühnen, P., and Stachelscheid, H. (2021). Generation of human induced pluripotent stem cell lines from 2 patients with MIRAGE syndrome. Stem Cell Res. 54, 102417. 10.1016/j.scr.2021.102417.

18. Vasimuddin, Md., Misra, S., Li, H., and Aluru, S. (2019). Efficient Architecture-Aware Acceleration of BWA-MEM for Multicore Systems. In 2019 IEEE International Parallel and Distributed Processing Symposium (IPDPS) (IEEE), pp. 314–324. 10.1109/IPDPS.2019.00041.

19. Babadi, M., Fu, J.M., Lee, S.K., Smirnov, A.N., Gauthier, L.D., Walker, M., Benjamin, D.I., Zhao, X., Karczewski, K.J., Wong, I., et al. (2023). GATK-gCNV enables discovery of rare copy number variants from exome sequencing data. Nat. Genet. 55, 1589–1597. 10.1038/s41588-023-01449-0.

20. Mölder, F., Jablonski, K.P., Letcher, B., Hall, M.B., Tomkins-Tinch, C.H., Sochat, V., Forster, J., Lee, S., Twardziok, S.O., Kanitz, A., et al. (2021). Sustainable data analysis with Snakemake. F1000Research 10, 33. 10.12688/f1000research.29032.2.

21. Mumm, S., Molini, B., Terrell, J., Srivastava, A., and Schlessinger, D. (1997). Evolutionary Features of the 4-Mb Xq21.3 XY Homology Region Revealed by a Map at 60-kb Resolution. Genome Res. 7, 307– 314. 10.1101/gr.7.4.307.

22. Ross, M.T., Grafham, D.V., Coffey, A.J., Scherer, S., McLay, K., Muzny, D., Platzer, M., Howell, G.R., Burrows, C., Bird, C.P., et al. (2005). The DNA sequence of the human X chromosome. Nature 434, 325–337. 10.1038/nature03440.

23. Webster, T.H., Couse, M., Grande, B.M., Karlins, E., Phung, T.N., Richmond, P.A., Whitford, W., and Wilson, M.A. (2019). Identifying, understanding, and correcting technical artifacts on the sex chromosomes in next-generation sequencing data. GigaScience 8, giz074. 10.1093/gigascience/giz074.

24. Diskin, S.J., Li, M., Hou, C., Yang, S., Glessner, J., Hakonarson, H., Bucan, M., Maris, J.M., and Wang, K. (2008). Adjustment of genomic waves in signal intensities from whole-genome SNP genotyping platforms. Nucleic Acids Res. 36, e126. 10.1093/nar/gkn556.

25. Wang, K., Li, M., Hadley, D., Liu, R., Glessner, J., Grant, S.F.A., Hakonarson, H., and Bucan, M. (2007). PennCNV: An integrated hidden Markov model designed for high-resolution copy number variation detection in whole-genome SNP genotyping data. Genome Res. 17, 1665–1674. 10.1101/gr.6861907.

26. DNAcopy Bioconductor. http://bioconductor.org/packages/DNAcopy/.

28. Martínez-Jiménez, F., Muiños, F., Sentís, I., Deu-Pons, J., Reyes-Salazar, I., Arnedo-Pac, C., Mularoni, L., Pich, O., Bonet, J., Kranas, H., et al. (2020). A compendium of mutational cancer driver genes. Nat. Rev. Cancer 20, 555–572. 10.1038/s41568-020-0290-x.

29. Illumina GenCall Data Analysis Software.

30. Pantazis, C.B., Yang, A., Lara, E., McDonough, J.A., Blauwendraat, C., Peng, L., Oguro, H., Kanaujiya, J., Zou, J., Sebesta, D., et al. (2022). A reference human induced pluripotent stem cell line for large-scale collaborative studies. Cell Stem Cell 29, 1685-1702.e22. 10.1016/j.stem.2022.11.004.

31. Gracia-Diaz, C., Perdomo, J.E., Khan, M.E., Roule, T., Disanza, B.L., Cajka, G.G., Lei, S., Gagne, A.L., Maguire, J.A., Shalem, O., et al. (2024). KOLF2.1J iPSCs carry CNVs associated with neurodevelopmental disorders. Cell Stem Cell 31, 288–289. 10.1016/j.stem.2024.02.007.

32. Ryan, M., McDonough, J.A., Ward, M.E., Cookson, M.R., Skarnes, W.C., and Merkle, F.T. (2024). Large structural variants in KOLF2.1J are unlikely to compromise neurological disease modeling. Cell Stem Cell 31, 290–291. 10.1016/j.stem.2024.02.006.

33. Fang, L., and Wang, K. (2018). Identification of Copy Number Variants from SNP Arrays Using PennCNV. In Copy Number Variants Methods in Molecular Biology., D. M. Bickhart, ed. (Springer New York), pp. 1–28. 10.1007/978-1-4939-8666-8_1.

34. Lavrichenko, K., Helgeland, Ø., Njølstad, P.R., Jonassen, I., and Johansson, S. (2021). SeeCiTe: a method to assess CNV calls from SNP arrays using trio data. Bioinformatics 37, 1876–1883. 10.1093/bioinformatics/btab028.

35. Zhang, Z., Cheng, H., Hong, X., Di Narzo, A.F., Franzen, O., Peng, S., Ruusalepp, A., Kovacic, J.C., Bjorkegren, J.L.M., Wang, X., et al. (2019). EnsembleCNV: an ensemble machine learning algorithm to identify and genotype copy number variation using SNP array data. Nucleic Acids Res. 47, e39–e39. 10.1093/nar/gkz068.

36. Gu, Z., and Mullighan, C.G. (2019). ShinyCNV: a Shiny/R application to view and annotate DNA copy number variations. Bioinformatics 35, 126–129. 10.1093/bioinformatics/bty546.

37. Zhang, X., Du, R., Li, S., Zhang, F., Jin, L., and Wang, H. (2014). Evaluation of copy number variation detection for a SNP array platform. BMC Bioinformatics 15, 1–9. 10.1186/1471-2105-15-50.

38. Dellinger, A.E., Saw, S.-M., Goh, L.K., Seielstad, M., Young, T.L., and Li, Y.-J. (2010). Comparative analyses of seven algorithms for copy number variant identification from single nucleotide polymorphism arrays. Nucleic Acids Res. 38, e105. 10.1093/nar/gkq040.

39. Van Loo, P., Nordgard, S.H., Lingjærde, O.C., Russnes, H.G., Rye, I.H., Sun, W., Weigman, V.J., Marynen, P., Zetterberg, A., Naume, B., et al. (2010). Allele-specific copy number analysis of tumors. Proc. Natl. Acad. Sci. 107, 16910–16915. 10.1073/pnas.1009843107.

40. Loh, P.-R., Genovese, G., Handsaker, R.E., Finucane, H.K., Reshef, Y.A., Palamara, P.F., Birmann, B.M., Talkowski, M.E., Bakhoum, S.F., McCarroll, S.A., et al. (2018). Insights into clonal haematopoiesis from 8,342 mosaic chromosomal alterations. Nature 559, 350–355. 10.1038/s41586-018-0321-x.

41. Loh, P.-R., Genovese, G., and McCarroll, S.A. (2020). Monogenic and polygenic inheritance become instruments for clonal selection. Nature 584, 136–141. 10.1038/s41586-020-2430-6.

42. Masood, D., Ren, L., Nguyen, C., Brundu, F.G., Zheng, L., Zhao, Y., Jaeger, E., Li, Y., Cha, S.W., Halpern, A., et al. (2024). Evaluation of somatic copy number variation detection by NGS technologies and bioinformatics tools on a hyper-diploid cancer genome. Genome Biol. 25, 163. 10.1186/s13059-024-03294-8.

43. Glessner, J.T., Chang, X., Liu, Y., Li, J., Khan, M., Wei, Z., Sleiman, P.M.A., and Hakonarson, H. (2021). MONTAGE: a new tool for high-throughput detection of mosaic copy number variation. BMC Genomics 22, 133. 10.1186/s12864-021-07395-7.

44. Wiegand, F., Lähnemann, D., Mölder, F., Uzuner, H., Prinz, A., Schramm, A., and Köster, J. (2025). Datavzrd: Rapid programming- and maintenance-free interactive visualization and communication of tabular data. PLOS ONE 20, e0323079. 10.1371/journal.pone.0323079.

