## Supplementary figures and images for "*StemCNV-check*: a pipeline for human pluripotent stem cell (hPSC) genomic integrity control using SNP array data and copy number variant scoring"

### Suplementary Figures

Von Kügelgen et al, Supplementary Figure 1

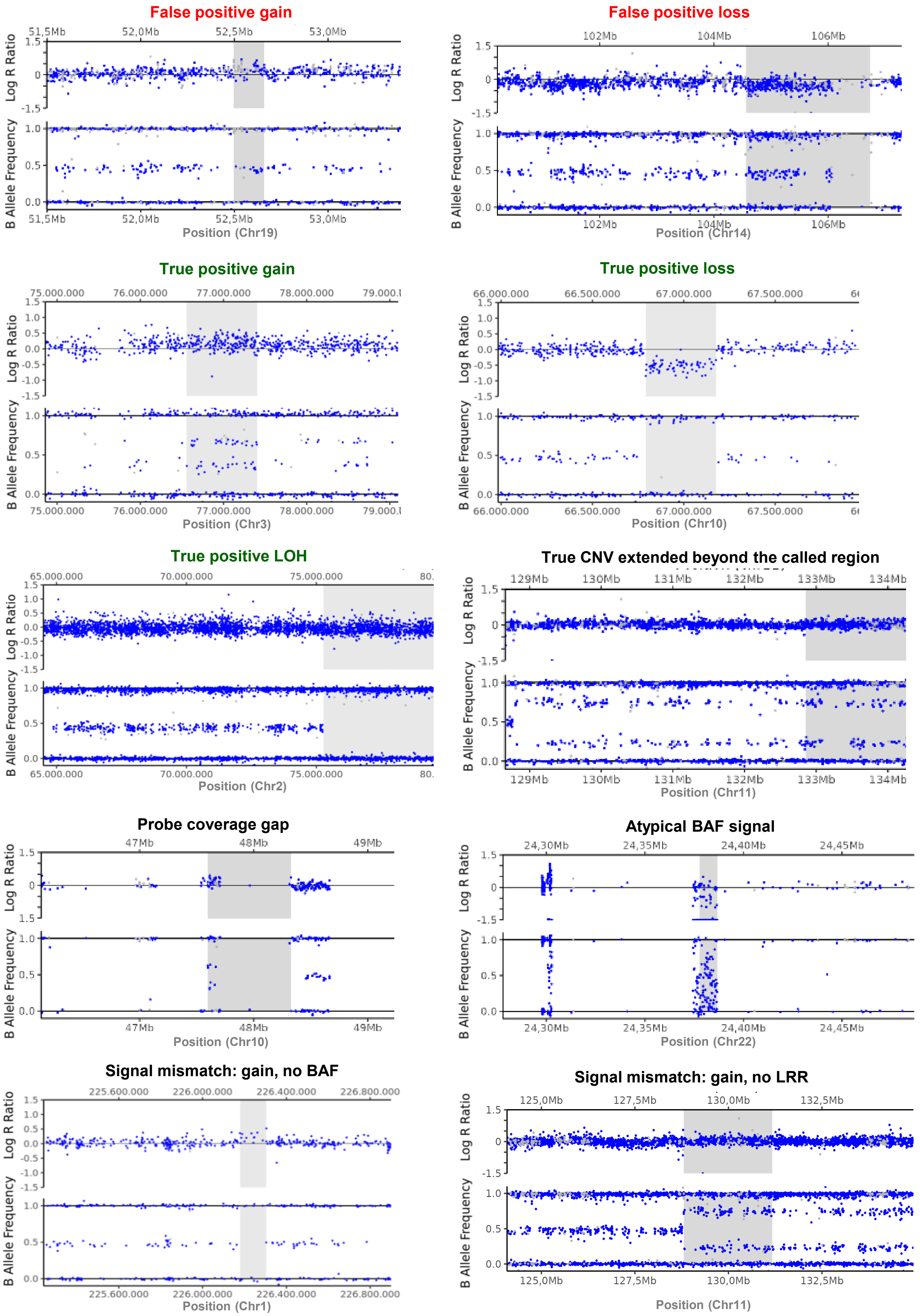

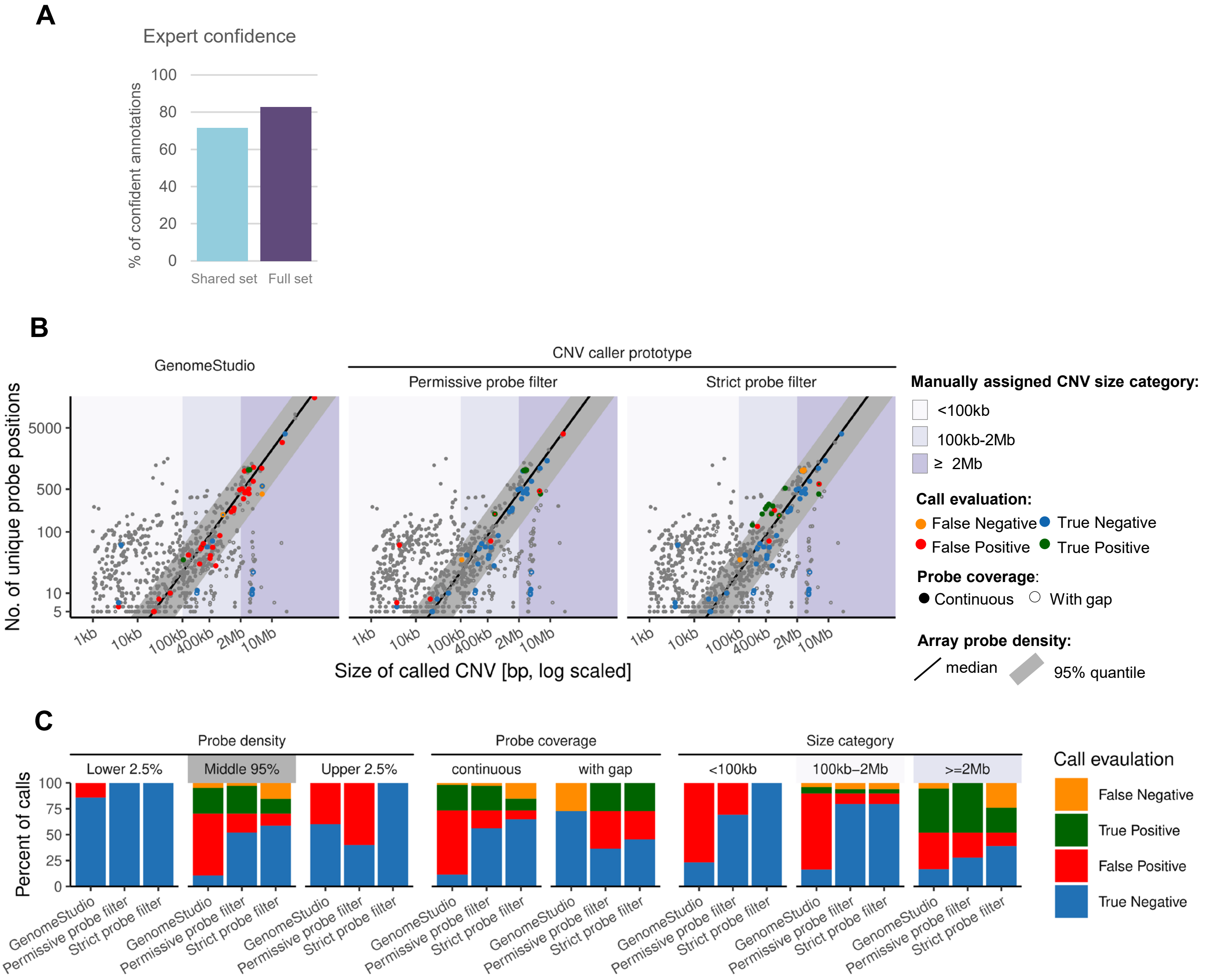

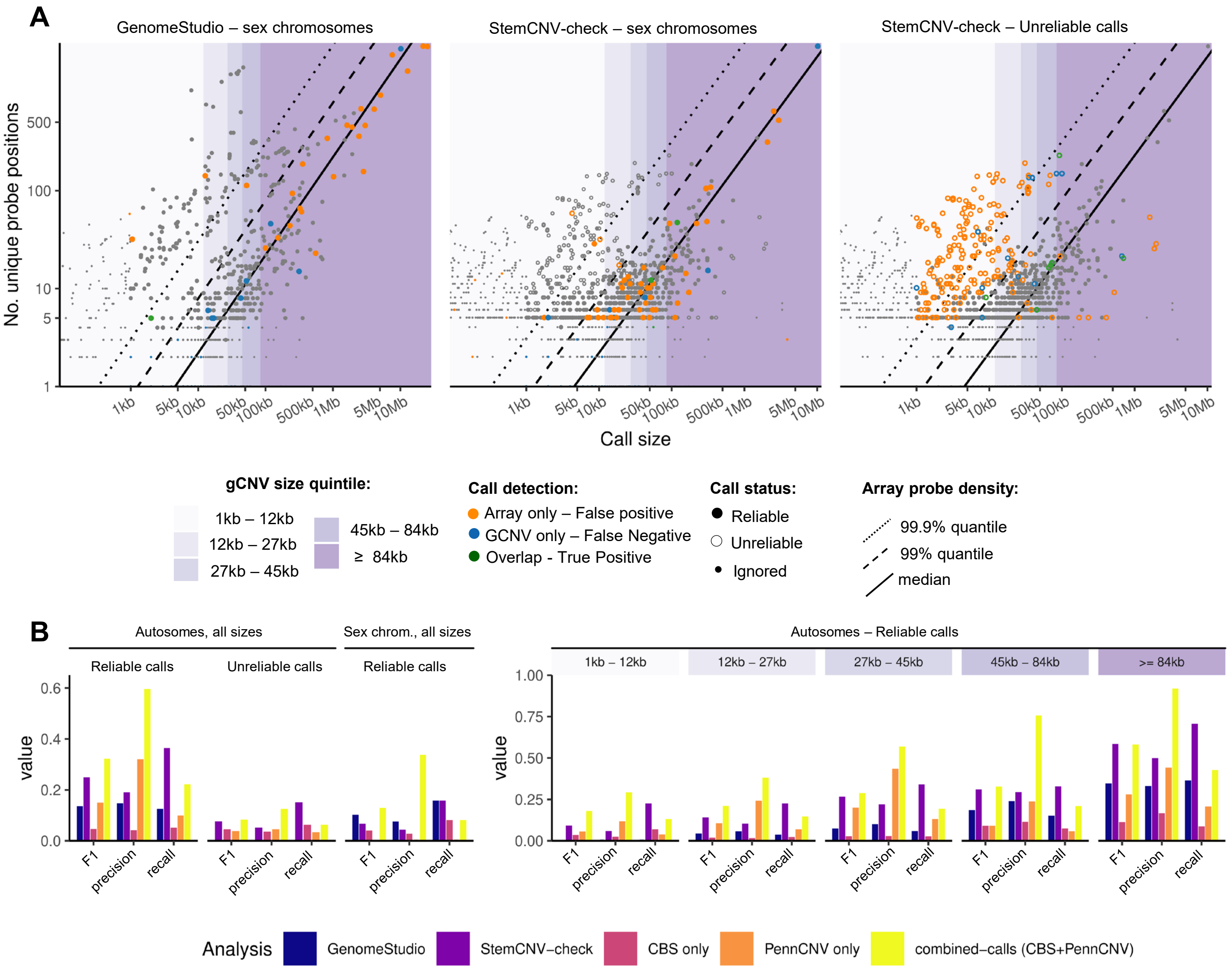

**Von Kügelgen et al, Supplementary Figure 4**

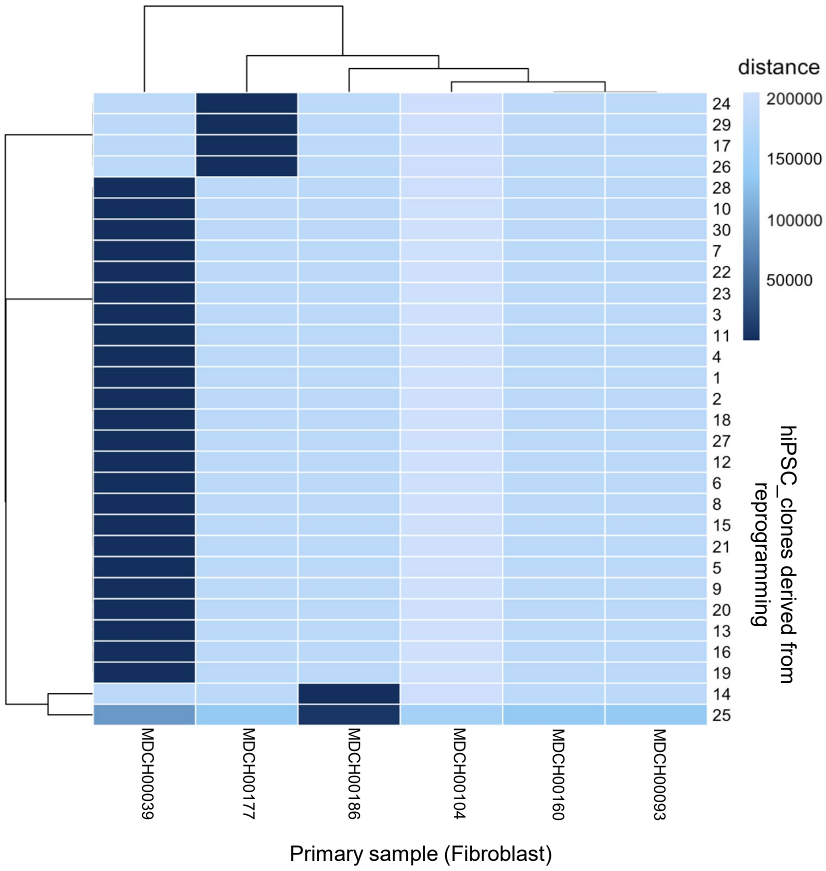

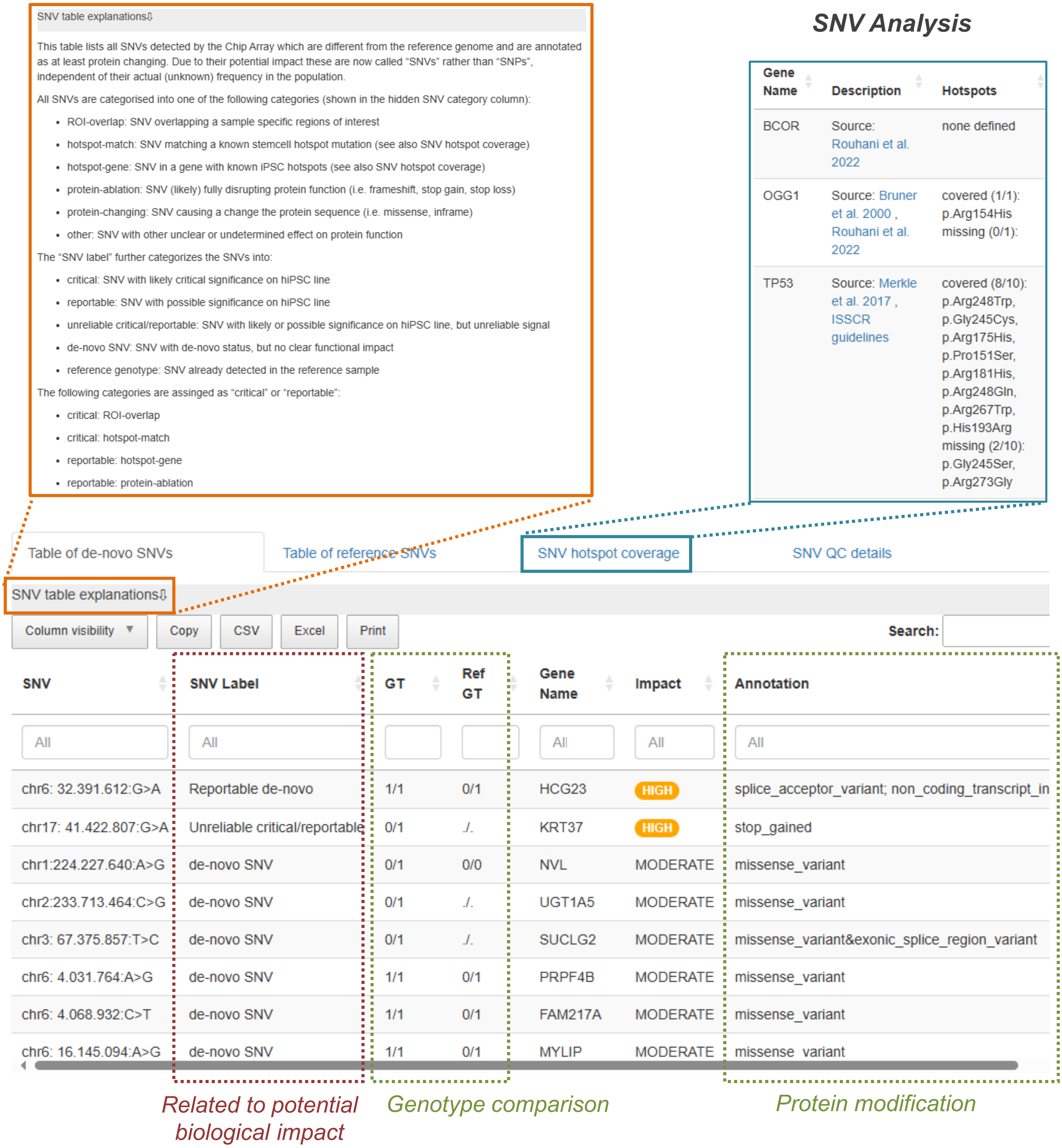

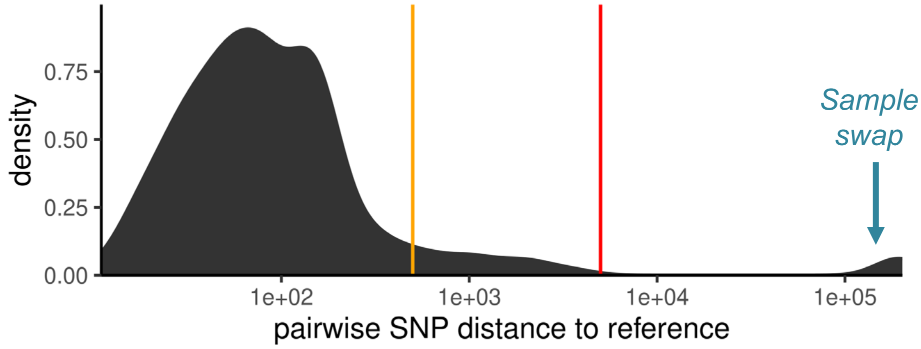
