## Supplementary Text for "*StemCNV-check*: a pipeline for human pluripotent stem cell (hPSC) genomic integrity control using SNP array data and copy number variant scoring"

### Supplementary Information

#### Supplementary Figures and Tables legends

**Supplementary Figure 1. Examples for different calls and their annotation in expert evaluation (related to Figure 1).**
Panels display array-derived log R ratio (LRR) and B-allele frequency (BAF) plots illustrating representative CNV calls from the evaluation dataset. Grey shading marks the CNV region.

**Supplementary Figure 2. Additional manual annotation summaries (related to Figure 1).**
**(A)** Bar plot presenting the proportion of confident expert annotations in the shared and full datasets. **(B)** Dot plots comparing CNV size (log scale) to number of unique probe positions for *GenomeStudio* and the CNV caller prototype under permissive and strict probe filter settings. Points are coloured by expert classification, grey points indicate autosomal CNVs, and hollow points indicate calls with a probe coverage gap. Background shading denotes size categories; diagonal line and grey band represent median probe density and middle 95% quantile. **(C)** Stacked bar plots summarizing the proportions of call classifications for sex chromosome CNVs, stratified by probe density, probe coverage type, and size category.

**Supplementary Figure 3. Performance in challenging regions and by individual callers (related to Figure 2).**
**(A)** Scatterplots of CNV size versus probe positions for *GenomeStudio* (sex chromosomes only), *StemCNV-check* (sex chromosomes only), and *StemCNV-check* (call status = *Unreliable*), with other calls shown in grey. Points are coloured by detection type and shaped by call status; background shading marks GCNV size quintiles. **(B)** Bar plots showing F1 score, precision, and recall for autosomes and sex chromosomes (*Reliable*/*Unreliable* calls), comparing *GenomeStudio*, *StemCNV-check*, CBS only, PennCNV only, and their intersection (combined calls).

**Supplementary Figure 4. SNP distance clustering of hiPSC clones and parental fibroblasts.**
Heatmap showing hierarchical clustering of pairwise SNP distances between hiPSC clones and their parental fibroblasts. Distance values are color-coded, with lighter shades indicating greater divergence. Intermediate shades displayed for clone 25 indicate its mixed genotype (no monoclonal).

**Supplementary Figure 5. SNV analysis and SNP distance distribution.**
**(A)** Interface screenshot of the SNV analysis section from the *StemCNV-check* report, showing table of de novo SNVs with annotation fields for gene name, genotype comparison, impact and functional effect. Insets in orange and blue show additional information which is hidden by default and becomes available by clicking the links in the interactive HTML report. **(B)** Density plot of pairwise SNP distances to reference samples from 261 array samples with a reference in the dataset. The x-axis is log-scaled; orange and red vertical lines mark default thresholds for warning (500) and high concern (5000). A distinct cluster at high distances indicates potential sample swaps.

**Supplementary Table 1. List of array and sequencing samples used in this study.**
Metadata for all SNP array-based and whole-genome sequencing (WGS) samples analysed. Columns include sample ID, collection source, sample group, sex, usage in individual figures, passage number, gene-editing status, type of modification, cell type, and array platform. For WGS data, median, standard deviation, and mean coverage, as well as number of mapped reads, are listed where available.

**Supplementary Table 2. Genome in a Bottle (GIAB) reference data used for benchmarking.**
List of FASTQ and paired FASTQ files from the NIST Genome in a Bottle (GIAB) reference dataset used for validation. Each entry includes file download links, MD5 checksums, NIST sample name, alias, and sex.

**Supplementary Table 3. SNP probe filter configurations tested in CNV calling.**
Summary of probe filtering strategies used for CNV calling, including workflow/tool name, probe filter name, number of probes per genomic position retained, treatment of pseudoautosomal and sex chromosome probes, GenCall (GC) and GenTrain (GT) thresholds, and figures where each configuration was applied.

**Supplementary Table 4. Curated CNV hotspot regions and annotations.**
List of recurrent CNV and SNV hotspot regions in human pluripotent stem cells (hPSCs), including genomic band location, call type (gain/loss), assigned *Check-Score* contribution, and associated gene content. Citations and DOIs for the original literature reporting each hotspot are provided below [1–40].

**Supplementary Table 5. Precision, recall, and F1 scores for CNV detection under varying conditions.**
Performance metrics for CNV calling across different probe filter settings, CNV callers, chromosome types, size ranges, and reliability flags. Columns include estimated precision, F1 score, recall, and a bracket description summarizing the parameter combination.

#### References

1. Närvä, E., Autio, R., Rahkonen, N., Kong, L., Harrison, N., Kitsberg, D., Borghese, L., Itskovitz-Eldor, J., Rasool, O., Dvorak, P., et al. (2010). High-resolution DNA analysis of human embryonic stem cell lines reveals culture-induced copy number changes and loss of heterozygosity. Nat. Biotechnol. *28*, 371–377. https://doi.org/10.1038/nbt.1615.

2. Kanchan, K., Iyer, K., Yanek, L.R., Carcamo-Orive, I., Taub, M.A., Malley, C., Baldwin, K., Becker, L.C., Broeckel, U., Cheng, L., et al. (2020). Genomic integrity of human induced pluripotent stem cells across nine studies in the NHLBI NextGen program. Stem Cell Res. *46*, 101803. https://doi.org/10.1016/j.scr.2020.101803.

3. Assou, S., Girault, N., Plinet, M., Bouckenheimer, J., Sansac, C., Combe, M., Mianné, J., Bourguignon, C., Fieldes, M., Ahmed, E., et al. (2020). Recurrent Genetic Abnormalities in Human Pluripotent Stem Cells: Definition and Routine Detection in Culture Supernatant by Targeted Droplet Digital PCR. Stem Cell Rep. *14*, 1–8. https://doi.org/10.1016/j.stemcr.2019.12.004.

4. Amps, K., Andrews, P.W., Anyfantis, G., Armstrong, L., Avery, S., Baharvand, H., Baker, J., Baker, D., Munoz, M.B., Beil, S., et al. (2011). Screening ethnically diverse human embryonic stem cells

identifies a chromosome 20 minimal amplicon conferring growth advantage. Nat. Biotechnol. *29*, 1132–1144. https://doi.org/10.1038/nbt.2051.

5. Tosca, L., Feraud, O., Magniez, A., Bas, C., Griscelli, F., Bennaceur-Griscelli, A., and Tachdjian, G. (2015). Genomic instability of human embryonic stem cell lines using different passaging culture methods. Mol. Cytogenet. *8*. https://doi.org/10.1186/s13039-015-0133-8.

6. Wu, H., Kim, K.J., Mehta, K., Paxia, S., Sundstrom, A., Anantharaman, T., Kuraishy, A.I., Doan, T., Ghosh, J., Pyle, A.D., et al. (2008). Copy Number Variant Analysis of Human Embryonic Stem Cells. Stem Cells *26*, 1484–1489. https://doi.org/10.1634/stemcells.2007-0993.

7. Baker, D., Hirst, A.J., Gokhale, P.J., Juarez, M.A., Williams, S., Wheeler, M., Bean, K., Allison, T.F., Moore, H.D., Andrews, P.W., et al. (2016). Detecting Genetic Mosaicism in Cultures of Human Pluripotent Stem Cells. Stem Cell Rep. *7*, 998–1012. https://doi.org/10.1016/j.stemcr.2016.10.003.

8. Martins-Taylor, K., Nisler, B.S., Taapken, S.M., Compton, T., Crandall, L., Montgomery, K.D., Lalande, M., and Xu, R.-H. (2011). Recurrent copy number variations in human induced pluripotent stem cells. Nat. Biotechnol. *29*, 488–491. https://doi.org/10.1038/nbt.1890.

9. Salomonis, N., Dexheimer, P.J., Omberg, L., Schroll, R., Bush, S., Huo, J., Schriml, L., Sui, S.H., Keddache, M., Mayhew, C., et al. (2016). Integrated Genomic Analysis of Diverse Induced Pluripotent Stem Cells from the Progenitor Cell Biology Consortium. Stem Cell Rep. *7*, 110–125. https://doi.org/10.1016/j.stemcr.2016.05.006.

10. Kang, X., Yu, Q., Huang, Y., Song, B., Chen, Y., Gao, X., He, W., Sun, X., and Fan, Y. (2015). Effects of Integrating and Non-Integrating Reprogramming Methods on Copy Number Variation and Genomic Stability of Human Induced Pluripotent Stem Cells. PLoS ONE *10*, e0131128. https://doi.org/10.1371/journal.pone.0131128.

11. Yamamoto, T., Sato, Y., Yasuda, S., Shikamura, M., Tamura, T., Takenaka, C., Takasu, N., Nomura, M., Dohi, H., Takahashi, M., et al. (2022). Correlation Between Genetic Abnormalities in Induced Pluripotent Stem Cell-Derivatives and Abnormal Tissue Formation in Tumorigenicity Tests. Stem Cells Transl. Med. *11*, 527–538. https://doi.org/10.1093/stcltm/szac014.

12. Merkle, F.T., Ghosh, S., Kamitaki, N., Mitchell, J., Avior, Y., Mello, C., Kashin, S., Mekhoubad, S., Ilic, D., Charlton, M., et al. (2017). Human pluripotent stem cells recurrently acquire and expand dominant negative P53 mutations. Nature *545*, 229–233. https://doi.org/10.1038/nature22312.

13. Kilpinen, H., Goncalves, A., Leha, A., Afzal, V., Alasoo, K., Ashford, S., Bala, S., Bensaddek, D., Casale, F.P., Culley, O.J., et al. (2017). Common genetic variation drives molecular heterogeneity in human iPSCs. Nature *546*, 370–375. https://doi.org/10.1038/nature22403.

14. Mus, L.M., Van Haver, S., Popovic, M., Trypsteen, W., Lefever, S., Zeltner, N., Ogando, Y., Jacobs, E.Z., Denecker, G., Sanders, E., et al. (2021). Recurrent chromosomal imbalances provide selective advantage to human embryonic stem cells under enhanced replicative stress conditions. Genes. Chromosomes Cancer *60*, 272–281. https://doi.org/10.1002/gcc.22931.

15. Lee, C.-T., Bendriem, R.M., Kindberg, A.A., Worden, L.T., Williams, M.P., Drgon, T., Mallon, B.S., Harvey, B.K., Richie, C.T., Hamilton, R.S., et al. (2015). Functional Consequences of 17q21.31/WNT3-WNT9B Amplification in hPSCs with Respect to Neural Differentiation. Cell Rep. *10*, 616–632. https://doi.org/10.1016/j.celrep.2014.12.050.

16. Amir, H., Touboul, T., Sabatini, K., Chhabra, D., Garitaonandia, I., Loring, J.F., Morey, R., and Laurent, L.C. (2017). Spontaneous Single-Copy Loss of TP53 in Human Embryonic Stem Cells Markedly Increases Cell Proliferation and Survival. Stem Cells *35*, 872–885. https://doi.org/10.1002/stem.2550.

17. Spits, C., Mateizel, I., Geens, M., Mertzanidou, A., Staessen, C., Vandeskelde, Y., Van der Elst, J., Liebaers, I., and Sermon, K. (2008). Recurrent chromosomal abnormalities in human embryonic stem cells. Nat. Biotechnol. *26*, 1361–1363. https://doi.org/10.1038/nbt.1510.

18. Werbowetski-Ogilvie, T.E., Bossé, M., Stewart, M., Schnerch, A., Ramos-Mejia, V., Rouleau, A., Wynder, T., Smith, M.-J., Dingwall, S., Carter, T., et al. (2009). Characterization of human embryonic stem cells with features of neoplastic progression. Nat. Biotechnol. *27*, 91–97. https://doi.org/10.1038/nbt.1516.

19. Avery, S., Hirst, A.J., Baker, D., Lim, C.Y., Alagaratnam, S., Skotheim, R.I., Lothe, R.A., Pera, M.F., Colman, A., Robson, P., et al. (2013). BCL-XL Mediates the Strong Selective Advantage of a 20q11.21 Amplification Commonly Found in Human Embryonic Stem Cell Cultures. Stem Cell Rep. *1*, 379–386. https://doi.org/10.1016/j.stemcr.2013.10.005.

20. Lefort, N., Feyeux, M., Bas, C., Féraud, O., Bennaceur-Griscelli, A., Tachdjian, G., Peschanski, M., and Perrier, A.L. (2008). Human embryonic stem cells reveal recurrent genomic instability at 20q11.21. Nat. Biotechnol. *26*, 1364–1366. https://doi.org/10.1038/nbt.1509.

21. Jo, H.-Y., Lee, Y., Ahn, H., Han, H.-J., Kwon, A., Kim, B.-Y., Ha, H.-Y., Kim, S.C., Kim, J.-H., Kim, Y.-O., et al. (2020). Functional in vivo and in vitro effects of 20q11.21 genetic aberrations on hPSC differentiation. Sci. Rep. *10*, 18582. https://doi.org/10.1038/s41598-020-75657-7.

22. Nguyen, H.T., Geens, M., Mertzanidou, A., Jacobs, K., Heirman, C., Breckpot, K., and Spits, C. (2014). Gain of 20q11.21 in human embryonic stem cells improves cell survival by increased expression of Bcl-xL. MHR Basic Sci. Reprod. Med. *20*, 168–177. https://doi.org/10.1093/molehr/gat077.

23. Zhang, J., Hirst, A.J., Duan, F., Qiu, H., Huang, R., Ji, Y., Bai, L., Zhang, F., Robinson, D., Jones, M., et al. (2019). Anti-apoptotic Mutations Desensitize Human Pluripotent Stem Cells to Mitotic Stress and Enable Aneuploid Cell Survival. Stem Cell Rep. *12*, 557–571. https://doi.org/10.1016/j.stemcr.2019.01.013.

24. Markouli, C., Couvreu De Deckersberg, E., Regin, M., Nguyen, H.T., Zambelli, F., Keller, A., Dziedzicka, D., De Kock, J., Tilleman, L., Van Nieuwerburgh, F., et al. (2019). Gain of 20q11.21 in Human Pluripotent Stem Cells Impairs TGF-β-Dependent Neuroectodermal Commitment. Stem Cell Rep. *13*, 163–176. https://doi.org/10.1016/j.stemcr.2019.05.005.

25. Na, J., Baker, D., Zhang, J., Andrews, P.W., and Barbaric, I. (2014). Aneuploidy in pluripotent stem cells and implications for cancerous transformation. Protein Cell *5*, 569–579. https://doi.org/10.1007/s13238-014-0073-9.

26. Mayshar, Y., Ben-David, U., Lavon, N., Biancotti, J.-C., Yakir, B., Clark, A.T., Plath, K., Lowry, W.E., and Benvenisty, N. (2010). Identification and Classification of Chromosomal Aberrations in Human Induced Pluripotent Stem Cells. Cell Stem Cell *7*, 521–531. https://doi.org/10.1016/j.stem.2010.07.017.

27. Herszfeld, D., Wolvetang, E., Langton-Bunker, E., Chung, T.-L., Filipczyk, A.A., Houssami, S., Jamshidi, P., Koh, K., Laslett, A.L., Michalska, A., et al. (2006). CD30 is a survival factor and a biomarker

for transformed human pluripotent stem cells. Nat. Biotechnol. *24*, 351–357. https://doi.org/10.1038/nbt1197.

28. Ben-David, U., Arad, G., Weissbein, U., Mandefro, B., Maimon, A., Golan-Lev, T., Narwani, K., Clark, A.T., Andrews, P.W., Benvenisty, N., et al. (2014). Aneuploidy induces profound changes in gene expression, proliferation and tumorigenicity of human pluripotent stem cells. Nat. Commun. *5*, 4825. https://doi.org/10.1038/ncomms5825.

29. Keller, A., Lei, Y., Krivec, N., De Deckersberg, E.C., Dziedzicka, D., Markouli, C., Sermon, K., Geens, M., and Spits, C. (2021). Gains of 12p13.31 delay WNT-mediated initiation of hPSC differentiation and promote residual pluripotency in a cell cycle dependent manner (Cell Biology) https://doi.org/10.1101/2021.05.22.445238.

30. Price, C.J., Stavish, D., Gokhale, P.J., Stevenson, B.A., Sargeant, S., Lacey, J., Rodriguez, T.A., and Barbaric, I. (2021). Genetically variant human pluripotent stem cells selectively eliminate wild-type counterparts through YAP-mediated cell competition. Dev. Cell *56*, 2455-2470.e10. https://doi.org/10.1016/j.devcel.2021.07.019.

31. Merkle, F.T., Ghosh, S., Genovese, G., Handsaker, R.E., Kashin, S., Meyer, D., Karczewski, K.J., O'Dushlaine, C., Pato, C., Pato, M., et al. (2022). Whole-genome analysis of human embryonic stem cells enables rational line selection based on genetic variation. Cell Stem Cell *29*, 472-486.e7. https://doi.org/10.1016/j.stem.2022.01.011.

32. Stavish, D., Price, C.J., Gelezauskaite, G., Alsehli, H., Leonhard, K.A., Taapken, S.M., McIntire, E.M., Laing, O., James, B.M., Riley, J.J., et al. (2024). Feeder-free culture of human pluripotent stem cells drives MDM4-mediated gain of chromosome 1q. Stem Cell Rep. *19*, 1217–1232. https://doi.org/10.1016/j.stemcr.2024.06.003.

33. Krivec, N., Couvreu De Deckersberg, E., Lei, Y., Al Delbany, D., Regin, M., Verhulst, S., Van Grunsven, L.A., Sermon, K., and Spits, C. (2024). Gain of 1q confers an MDM4-driven growth advantage to undifferentiated and differentiating hESC while altering their differentiation capacity. Cell Death Dis. *15*, 852. https://doi.org/10.1038/s41419-024-07236-x.

34. Stavish, D., Price, C.J., Gelezauskaite, G., Leonhard, K.A., Taapken, S.M., McIntire, E.M., Laing, O., James, B.M., Riley, J.J., Zerbib, J., et al. (2023). Cytogenetic resource enables mechanistic resolution of changing trends in human pluripotent stem cell aberrations linked to feeder-free culture. Preprint at bioRxiv, https://doi.org/10.1101/2023.09.21.558777 https://doi.org/10.1101/2023.09.21.558777.

35. Halliwell, J., Barbaric, I., and Andrews, P.W. (2020). Acquired genetic changes in human pluripotent stem cells: origins and consequences. Nat. Rev. Mol. Cell Biol. *21*, 715–728. https://doi.org/10.1038/s41580-020-00292-z.

36. Lei, Y., Al Delbany, D., Krivec, N., Regin, M., Couvreu De Deckersberg, E., Janssens, C., Ghosh, M., Sermon, K., and Spits, C. (2024). SALL3 mediates the loss of neuroectodermal differentiation potential in human embryonic stem cells with chromosome 18q loss. Stem Cell Rep. *19*, 562–578. https://doi.org/10.1016/j.stemcr.2024.03.001.

37. Gore, A., Li, Z., Fung, H.-L., Young, J.E., Agarwal, S., Antosiewicz-Bourget, J., Canto, I., Giorgetti, A., Israel, M.A., Kiskinis, E., et al. (2011). Somatic coding mutations in human induced pluripotent stem cells. Nature *471*, 63–67. https://doi.org/10.1038/nature09805.

38. Rouhani, F.J., Zou, X., Danecek, P., Badja, C., Amarante, T.D., Koh, G., Wu, Q., Memari, Y., Durbin, R., Martincorena, I., et al. (2022). Substantial somatic genomic variation and selection for BCOR mutations in human induced pluripotent stem cells. Nat. Genet. *54*, 1406–1416. https://doi.org/10.1038/s41588-022-01147-3.

39. Avior, Y., Lezmi, E., Eggan, K., and Benvenisty, N. (2021). Cancer-Related Mutations Identified in Primed Human Pluripotent Stem Cells. Cell Stem Cell *28*, 10–11. https://doi.org/10.1016/j.stem.2020.11.013.

40. Bruner, S.D., Norman, D.P.G., and Verdine, G.L. (2000). Structural basis for recognition and repair of the endogenous mutagen 8-oxoguanine in DNA. Nature *403*, 859–866. https://doi.org/10.1038/35002510.
