## Supplementary material for "*StemCNV-check*: a pipeline for human pluripotent stem cell (hPSC) genomic integrity control using SNP array data and copy number variant scoring": Example report: StemCNV-check-_example-report.html

BIHi001-B\_MB01 StemCNV check report


### BIHi001-B\_MB01 StemCNV check report

###### StemCNV-check, version: 1.0.0

### Sample Overview

#### Sample Information

#### QC metrics

##### Summary

Data QC explanations

⇩

These summary tables are meant to serve as a quick overview of the
quality of an hPSC sample. This first table contains QC metrics
primarily related to the SNP data quality (affected by both the DNA used
and the array run itself), this table will also display values from the
reference sample if possible. The second table contains QC metrics
related to the potentially problematic CNVs and SNVs identified in only
the analysed sample.

Coloring of all fields is based on (usually) two thresholds defined in
the config file (under the evaluation\_settings section), one for a
signalling (yellow) level, and one for a more serious warning (orange)
or even critical (red) level. Only certain values are potentially
considered critical and are marked by bold text in the table, which
values behave like this is also defined in the config.


Sample QC explanations

⇩

This table sums up all variant findings from the analysed sample,
which were flagged as critical or reportable.

Note that in contrast to general SNP probes on the array, only those
single variants that actually show an alternative allele and affect a
protein are considered SNVs by StemCNV-check. Variants that match the
genotype of assigned reference sample are never considered critical or
reportable.

The following criteria are used to assign SNVs as critical or
reportable:

Critical SNVs:

- hotspot-match: SNV matching a known stemcell hotspot mutation (see
  also SNV hotspot coverage)

Reportable SNVs:

- hotspot-gene: SNV in a gene with known iPSC hotspots (see also
  SNV hotspot coverage)
- protein-ablation: SNV (likely) fully disrupting protein function
  (i.e. frameshift, stop gain, stop loss)

For copy number variants (CNVs) the assigned label designation takes
into account a minimum Check\_Score threshold, overlap with a reference
call and certain call filter flags (see below). The defined call label
criteria are:

- Critical de-novo:

  Minimum required Check-Score: 55

  Exclusion of calls with any filter among: high\_probe\_dens, probe\_gap,
  min\_size, min\_probes, min\_density

  Match to CNV in reference sample: not allowed
- Reportable de-novo:

  Minimum required Check-Score: 55

  Exclusion of calls with any filter among: min\_size, min\_probes,
  min\_density

  Match to CNV in reference sample: not allowed
- de-novo call:

  Minimum required Check-Score: 0

  Exclusion of calls with any filter among: min\_size, min\_probes,
  min\_density

  Match to CNV in reference sample: not allowed
- Reference genotype:

  Minimum required Check-Score: 0

  Exclusion of calls with any filter among:

  Match to CNV in reference sample: required
- Excluded call:

  Minimum required Check-Score: 0

  Exclusion of calls with any filter among:

  Match to CNV in reference sample: not allowed

The defined CNV filter flags are:

- min\_size: CNV call below minimum size (<1000bp)
- min\_probes: CNV call from <5 probes
- min\_density: CNV call with <10 probes/Mb
- high\_probe\_dens: Probe density of segment is higher than 99% of
  the array
- probe\_gap: Probe coverage of segment has considerable gap (min.
  33% depending on probe number - see config)

##### GenCall

This table displays the direct quality metrics from the GenCall
software.

##### PennCNV

The first table displays quality metrics from the PennCNV algorythm.
The second displays CNV call statistcs for only PennCNV

##### CBS

This table displays CNV call statistcs for only CBS.

##### Config

Changes from default config:

```
values_changed:
  'config: settings : SNV_analysis : SNP_clustering : max_number_samples':
    new_value: 25
    old_value: 20
iterable_item_added:
  'config: settings : SNV_analysis : SNP_clustering : sample_ids[0]': CCD1112Sk_HFF
  'config: settings : SNV_analysis : SNP_clustering : sample_ids[1]': BIHi001-A_MB01
  'config: settings : SNV_analysis : SNP_clustering : sample_ids[2]': BIHi001-A_WB03
  'config: settings : SNV_analysis : SNP_clustering : sample_ids[3]': BIHi001-A_WB04
  'config: settings : SNV_analysis : SNP_clustering : sample_ids[4]': BIHi001-B_MB01
  'config: settings : SNV_analysis : SNP_clustering : sample_ids[5]': BIHi001-B_WB04
  'config: settings : SNV_analysis : SNP_clustering : sample_ids[6]': SCVI_111
  'config: settings : SNV_analysis : SNP_clustering : sample_ids[7]': BIHi005-A_MB02_1
  'config: settings : SNV_analysis : SNP_clustering : sample_ids[8]': BIHi005-A_MB02_2
  'config: settings : SNV_analysis : SNP_clustering : sample_ids[9]': BIHi005-A_MB02_3
  'config: settings : SNV_analysis : SNP_clustering : sample_ids[10]': BIHi005-A_WB02
  'config: settings : SNV_analysis : SNP_clustering : sample_ids[11]': BIHi005-A_WB04
  'config: settings : SNV_analysis : SNP_clustering : sample_ids[12]': NHDF_lot0000477954
  'config: settings : SNV_analysis : SNP_clustering : sample_ids[13]': BIHi250-A_MB01
  'config: settings : SNV_analysis : SNP_clustering : sample_ids[14]': BIHi250-A_WB01
  'config: settings : SNV_analysis : SNP_clustering : sample_ids[15]': BIHi250-A_WB02
  'config: settings : SNV_analysis : SNP_clustering : sample_ids[16]': BIHi250-A_WB01_2
  'config: settings : SNV_analysis : SNP_clustering : sample_ids[17]': BIHi250-A_WB03
  'config: settings : SNV_analysis : SNP_clustering : sample_ids[18]': KOLF21J
  'config: settings : SNV_analysis : SNP_clustering : sample_ids[19]': UCSFi001-A_MB01
  'config: settings : SNV_analysis : SNP_clustering : sample_ids[20]': UCSFi001-A_WB01
```

Complete config used by StemCNV-check:

```
array_definition:
  GSAMD-24v3-hg38:
    genome_version: hg38
    bpm_manifest_file: ../static-data/GSAMD-24v3-0-EA_20034606_A2.bpm
    egt_cluster_file: ../static-data/GSAMD-24v3-0-EA_20034606_A1.egt
    csv_manifest_file: ../static-data/GSAMD-24v3-0-EA_20034606_A2.csv
    penncnv_GCmodel_file: ../static-data/PennCNV-GCmodel_hg38_GSAMD-v24.gcmodel
    array_density_file: ../static-data/density_hg38_GSAMD-v24.bed
    array_gaps_file: ../static-data/gaps_hg38_GSAMD-v24.bed
    penncnv_pfb_file: ../static-data/PennCNV-PFB_hg38_GSAMD-v24.pfb
raw_data_folder: ../RAW_DATA
data_path: data_reports
log_path: data_reports/logs
evaluation_settings:
  CNV_call_labels:
    Critical de-novo:
      minimum_check_score: 55
      not_allowed_vcf_filters:
      - high_probe_dens
      - probe_gap
      - min_size
      - min_probes
      - min_density
      reference_match: no
    Reportable de-novo:
      minimum_check_score: 55
      not_allowed_vcf_filters:
      - min_size
      - min_probes
      - min_density
      reference_match: no
    de-novo call:
      minimum_check_score: 0
      not_allowed_vcf_filters:
      - min_size
      - min_probes
      - min_density
      reference_match: no
    Reference genotype:
      minimum_check_score: 0
      not_allowed_vcf_filters: ~
      reference_match: yes
    Excluded call:
      minimum_check_score: 0
      not_allowed_vcf_filters: ~
      reference_match: no
  summary_stat_warning_levels:
    call_rate:
    - 0.99
    - 0.99
    SNP_pairwise_distance_to_reference:
    - 500
    - 5000
    loss_gain_log2ratio:
    - 2
    - 4
    total_calls_CNV:
    - 10
    - 50
    total_calls_LOH:
    - 30
    - 75
    reportable_calls_CNV:
    - 5
    - 10
    reportable_calls_LOH:
    - 5
    - 10
    critical_calls_CNV:
    - 1
    - 1
    critical_calls_LOH:
    - 1
    - 1
    reportable_SNVs:
    - 5
    - 10
    critical_SNVs:
    - 1
    - 1
    call_count_excl_labels: Excluded call
    use_last_level:
    - call_rate
    - computed_gender
    - SNP_pairwise_distance_to_reference
    - critical_SNVs
    - critical_calls_CNV
    - critical_calls_LOH
  collate_output:
    file_format: xlsx
    summary_extra_sampletable_cols: Reference_Sample
    cnv_collate_call_selection:
      whitelist_call_label: ~
      blacklist_call_label: Excluded call
global_settings:
  cache_dir: ~/work/.stem-cnv-check
  hg19_mehari_transcript_db: ../static-data/mehari-data-txs-GRCh37-ensembl-0.10.3.bin.zst
  hg38_mehari_transcript_db: ../static-data/mehari-data-txs-GRCh38-ensembl-0.10.3.bin.zst
  dosage_sensitivity_scores: __cache-default__
  hg19_genome_fasta: /data/cephfs-1/work/projects/cubit/current/static_data/reference/GRCh37/hs37d5/hs37d5.fa
  hg38_genome_fasta: /data/cephfs-1/work/projects/cubit/current/static_data/reference/GRCh38/hs38/hs38.fa
  hg19_gtf_file: /data/cephfs-1/work/projects/cubit/current/static_data/annotation/GENCODE/19/GRCh37/gencode.v19.annotation.gtf
  hg38_gtf_file: /data/cephfs-1/work/projects/cubit/current/static_data/annotation/GENCODE/33/GRCh38/gencode.v33.annotation.gtf
  hg19_genomeInfo_file: ../static-data/UCSC_hg19_chromosome-info.tsv
  hg38_genomeInfo_file: ../static-data/UCSC_hg38_chromosome-info.tsv
settings:
  CNV.calling.tools:
  - PennCNV
  - CBS
  probe_filter_sets:
    standard:
      GenTrainScore: 0.15
      GenCallScore: 0.15
      Position.duplicates: highest-GenCall
      Pseudoautosomal: remove-male
  default_probe_filter_set: standard
  PennCNV:
    probe_filter_settings: _default_
    enable_LOH_calls: yes
    call.merging:
      merge.gap.absolute: 500
      merge.gap.snps: 10
      call.extension.percent: 60
      maximum.gap.allowed: 500000
    filter.minprobes: 5
    filter.minlength: 1000
    filter.mindensity.Mb: 10
  CBS:
    probe_filter_settings: _default_
    undo.SD.val: 1
    call.merging:
      merge.gap.absolute: 500
      merge.gap.snps: 10
      call.extension.percent: 60
      maximum.gap.allowed: 500000
    filter.minprobes: 5
    filter.minlength: 1000
    filter.mindensity.Mb: 10
    LRR.loss: -0.25
    LRR.loss.large: -1.1
    LRR.gain: 0.2
    LRR.gain.large: 0.75
    LRR.male.XorY.loss: -0.5
    LRR.male.XorY.gain: 0.28
    LRR.male.XorY.gain.large: 0.75
    LRR.female.X.loss: -0.05
    LRR.female.XX.loss: -0.9
    LRR.female.X.gain: 0.5
    LRR.female.X.gain.large: 1.05
  array_attribute_summary:
    density.windows: 100000
    min.gap.size: auto-array
  CNV_processing:
    call_processing:
      probe_filter_settings: _default_
      tool.overlap.greatest.call.min.perc: 50
      tool.overlap.min.cov.sum.perc: 60
      filter.minprobes: 5
      filter.minlength: 1000
      filter.mindensity.Mb: 10
      min.reciprocal.coverage.with.ref: 50
      gap_area.uniq_probes.rel:
      - -12.0
      - 12.5
      min.perc.gap_area: 0.33
      density.quantile.cutoff: 0.99
    gene_overlap:
      exclude_gene_type_regex: ~
      include_only_these_gene_types:
      - lncRNA
      - miRNA
      - protein_coding
      whitelist_hotspot_genes: yes
      stemcell_hotspot_list: __inbuilt__/supplemental-files/CNV-stemcell-hotspots.tsv
      cancer_gene_list: __inbuilt__/supplemental-files/genelist-cancer-drivers.tsv
      dosage_sensitive_gene_name_fixes: __inbuilt__/supplemental-files/gene-names-mapping-dosage-sensitivity.tsv
    Check_score_values:
      pHaplo_threshold: 0.86
      pTriplo_threshold: 0.94
      dosage_sensitive_gene: 5
      any_other_gene: 0.2
      single_copy_factor: 0.333
      double_copy_factor: 0.5
      neutral_copy_factor: 0.275
      flat_decrease: 15
    precision_estimation_file: __inbuilt__/supplemental-files/precision_estimates.tsv
  SNV_analysis:
    probe_filter_settings: _default_
    snv_hotspot_table: __inbuilt__/supplemental-files/SNV-stemcell-hotspots.tsv
    flag_GenCall_minimum: 0.2
    variant_selection:
      Impact:
      - HIGH
      - MODERATE
      Annotation_regex: ~
      include_all_ROI_overlaps: yes
    critical_SNV: hotspot-match
    reportable_SNV:
    - hotspot-gene
    - protein-ablation
    protein_ablation_annotations:
      Impact: HIGH
      Annotation_regex: ~
    protein_change_annotations:
      Impact: ~
      Annotation_regex: missense_variant|inframe
    SNP_clustering:
      sample_ids:
      - CCD1112Sk_HFF
      - BIHi001-A_MB01
      - BIHi001-A_WB03
      - BIHi001-A_WB04
      - BIHi001-B_MB01
      - BIHi001-B_WB04
      - SCVI_111
      - BIHi005-A_MB02_1
      - BIHi005-A_MB02_2
      - BIHi005-A_MB02_3
      - BIHi005-A_WB02
      - BIHi005-A_WB04
      - NHDF_lot0000477954
      - BIHi250-A_MB01
      - BIHi250-A_WB01
      - BIHi250-A_WB02
      - BIHi250-A_WB01_2
      - BIHi250-A_WB03
      - KOLF21J
      - UCSFi001-A_MB01
      - UCSFi001-A_WB01
      id_columns: ~
      match_columns:
      - Chip_Name
      - Sample_Group
      max_number_samples: 25
  vcf_output:
    chrom_style: UCSC
reports:
  StemCNV-check-report:
    file_type: html
  _default_:
    include_sections: __all__
    exclude_sections: ~
    sample.info.extra.cols:
    - Chip_Name
    - Chip_Pos
    CNV_call_labels_removed: Excluded call
    call.data.and.plots:
      _default_:
        min_number_plots: 20
        always_include_CNVs: ~
        include.plot: yes
        include.hotspot.table: yes
        include.gene.table.details: Call
        plot.flanking.region.relative: 2
        plot.region.minsize: 2000000
      denovo:
        min_number_plots: 20
        always_include_CNVs:
        - Critical de-novo
        - Reportable de-novo
        include.plot: yes
        include.hotspot.table: yes
        include.gene.table.details: Call
        plot.flanking.region.relative: 2
        plot.region.minsize: 2000000
        call_labels_include:
        - Critical de-novo
        - Reportable de-novo
        - de-novo call
      reference_gt:
        min_number_plots: 20
        always_include_CNVs: ~
        include.plot: yes
        include.hotspot.table: yes
        include.gene.table.details: Call
        plot.flanking.region.relative: 2
        plot.region.minsize: 2000000
        call_labels_include: Reference genotype
      regions_of_interest:
        min_number_plots: 20
        always_include_CNVs: ~
        include.plot: yes
        include.hotspot.table: yes
        include.gene.table.details: Call
        plot.flanking.region.relative: 2
        plot.region.minsize: 100000
    SNP_comparison:
      dendrogram.color.by: Sample_Group
      dendrogram.shape.by: Chip_Name
    genome_overview:
      call_labels_overview:
      - Critical de-novo
      - Reportable de-novo
      - de-novo call
      - Reference genotype
      show_reference: yes
wildcard_constraints:
  sample_id: '[a-zA-Z0-9-_]+'
  sentrix_pos: R[0-9]{2}C[0-9]{2}
  sentrix_name: '[0-9]+'
tools:
  _default_:
    threads: 1
    memory: 6000
    runtime: 2h
    partition: medium
  GenCall:
    threads: 4
    memory: 8000
    runtime: 4h
  CBS:
    memory: 6000
    runtime: 1h
  CNV.process:
    memory: 6000
    runtime: 1h
  PennCNV:
    memory: 4000
    runtime: 1h
  SNV_analysis:
    threads: 2
    memory: 20000
    runtime: 4h
  knitr:
    memory: 25000
    runtime: 1h
  gtc2vcf:
    memory: 6000
sample_table: sample_table_reports.xlsx
column_remove_regex: None
basedir: /data/cephfs-1/work/projects/stachelscheid-cellline-arrays/manusscript_2024
configfile: config_reports.yaml
target: complete
cache_path: /data/cephfs-1/home/users/vonkunic_c/work/.stem-cnv-check
verbose_level: 0
is_wsl: 0
snakedir: /data/cephfs-1/work/groups/cubi/users/vonkunic_c/git-repos/StemCNV-check/src/stemcnv_check
report_settings:
  include_sections: __all__
  exclude_sections: ~
  sample.info.extra.cols:
  - Chip_Name
  - Chip_Pos
  CNV_call_labels_removed: Excluded call
  call.data.and.plots:
    _default_:
      min_number_plots: 20
      always_include_CNVs: ~
      include.plot: yes
      include.hotspot.table: yes
      include.gene.table.details: Call
      plot.flanking.region.relative: 2
      plot.region.minsize: 2000000
    denovo:
      min_number_plots: 20
      always_include_CNVs:
      - Critical de-novo
      - Reportable de-novo
      include.plot: yes
      include.hotspot.table: yes
      include.gene.table.details: Call
      plot.flanking.region.relative: 2
      plot.region.minsize: 2000000
      call_labels_include:
      - Critical de-novo
      - Reportable de-novo
      - de-novo call
    reference_gt:
      min_number_plots: 20
      always_include_CNVs: ~
      include.plot: yes
      include.hotspot.table: yes
      include.gene.table.details: Call
      plot.flanking.region.relative: 2
      plot.region.minsize: 2000000
      call_labels_include: Reference genotype
    regions_of_interest:
      min_number_plots: 20
      always_include_CNVs: ~
      include.plot: yes
      include.hotspot.table: yes
      include.gene.table.details: Call
      plot.flanking.region.relative: 2
      plot.region.minsize: 100000
  SNP_comparison:
    dendrogram.color.by: Sample_Group
    dendrogram.shape.by: Chip_Name
  genome_overview:
    call_labels_overview:
    - Critical de-novo
    - Reportable de-novo
    - de-novo call
    - Reference genotype
    show_reference: yes
  file_type: html
```

##### R session info

```
## ─ Session info ───────────────────────────────────────────────────────────────
##  setting  value
##  version  R version 4.3.3 (2024-02-29)
##  os       Rocky Linux 9.6 (Blue Onyx)
##  system   x86_64, linux-gnu
##  ui       X11
##  language (EN)
##  collate  C.UTF-8
##  ctype    C.UTF-8
##  tz       Europe/Berlin
##  date     2025-09-23
##  pandoc   3.6.4 @ /data/cephfs-1/home/users/vonkunic_c/work/.stem-cnv-check/f4f9fb2443eadf59b8a94b12a19c1c0f_/bin/ (via rmarkdown)
##  quarto   NA
## 
## ─ Packages ───────────────────────────────────────────────────────────────────
##  package              * version    date (UTC) lib source
##  abind                  1.4-5      2016-07-21 [1] CRAN (R 4.3.3)
##  ape                    5.8-1      2024-12-16 [1] CRAN (R 4.3.3)
##  backports              1.5.0      2024-05-23 [1] CRAN (R 4.3.3)
##  base64enc              0.1-3      2015-07-28 [1] CRAN (R 4.3.3)
##  Biobase                2.62.0     2023-10-24 [1] Bioconductor
##  BiocGenerics         * 0.48.1     2023-11-01 [1] Bioconductor
##  BiocIO                 1.12.0     2023-10-24 [1] Bioconductor
##  BiocParallel           1.36.0     2023-10-24 [1] Bioconductor
##  Biostrings             2.70.1     2023-10-25 [1] Bioconductor
##  bit                    4.6.0      2025-03-06 [1] CRAN (R 4.3.3)
##  bit64                  4.6.0-1    2025-01-16 [1] CRAN (R 4.3.3)
##  bitops                 1.0-9      2024-10-03 [1] CRAN (R 4.3.3)
##  broom                  1.0.8      2025-03-28 [1] CRAN (R 4.3.3)
##  bslib                  0.9.0      2025-01-30 [1] CRAN (R 4.3.3)
##  cachem                 1.1.0      2024-05-16 [1] CRAN (R 4.3.3)
##  car                    3.1-3      2024-09-27 [1] CRAN (R 4.3.3)
##  carData                3.0-5      2022-01-06 [1] CRAN (R 4.3.3)
##  cellranger             1.1.0      2016-07-27 [1] CRAN (R 4.3.3)
##  cli                    3.6.4      2025-02-13 [1] CRAN (R 4.3.3)
##  cluster                2.1.8.1    2025-03-12 [1] CRAN (R 4.3.3)
##  codetools              0.2-20     2024-03-31 [1] CRAN (R 4.3.3)
##  colorspace             2.1-1      2024-07-26 [1] CRAN (R 4.3.3)
##  crayon                 1.5.3      2024-06-20 [1] CRAN (R 4.3.3)
##  crosstalk              1.2.1      2023-11-23 [1] CRAN (R 4.3.3)
##  DelayedArray           0.28.0     2023-10-24 [1] Bioconductor
##  dendextend           * 1.19.0     2024-11-15 [1] CRAN (R 4.3.3)
##  digest                 0.6.37     2024-08-19 [1] CRAN (R 4.3.3)
##  dplyr                * 1.1.4      2023-11-17 [1] CRAN (R 4.3.3)
##  DT                   * 0.33       2024-04-04 [1] CRAN (R 4.3.3)
##  evaluate               1.0.3      2025-01-10 [1] CRAN (R 4.3.3)
##  farver                 2.1.2      2024-05-13 [1] CRAN (R 4.3.3)
##  fastmap                1.2.0      2024-05-15 [1] CRAN (R 4.3.3)
##  forcats              * 1.0.0      2023-01-29 [1] CRAN (R 4.3.3)
##  Formula                1.2-5      2023-02-24 [1] CRAN (R 4.3.3)
##  generics               0.1.3      2022-07-05 [1] CRAN (R 4.3.3)
##  GenomeInfoDb         * 1.38.1     2023-11-08 [1] Bioconductor
##  GenomeInfoDbData       1.2.11     2025-04-03 [1] Bioconductor
##  GenomicAlignments      1.38.0     2023-10-24 [1] Bioconductor
##  GenomicRanges        * 1.54.1     2023-10-29 [1] Bioconductor
##  ggplot2              * 3.5.1      2024-04-23 [1] CRAN (R 4.3.3)
##  ggpubr               * 0.6.0      2023-02-10 [1] CRAN (R 4.3.3)
##  ggrepel              * 0.9.6      2024-09-07 [1] CRAN (R 4.3.3)
##  ggsignif               0.6.4      2022-10-13 [1] CRAN (R 4.3.3)
##  glue                   1.8.0      2024-09-30 [1] CRAN (R 4.3.3)
##  gridExtra              2.3        2017-09-09 [1] CRAN (R 4.3.3)
##  grImport2              0.3-3      2024-07-30 [1] CRAN (R 4.3.3)
##  gtable                 0.3.6      2024-10-25 [1] CRAN (R 4.3.3)
##  hms                    1.1.3      2023-03-21 [1] CRAN (R 4.3.3)
##  htmltools              0.5.8.1    2024-04-04 [1] CRAN (R 4.3.3)
##  htmlwidgets            1.6.4      2023-12-06 [1] CRAN (R 4.3.3)
##  IRanges              * 2.36.0     2023-10-24 [1] Bioconductor
##  jpeg                   0.1-11     2025-03-21 [1] CRAN (R 4.3.3)
##  jquerylib              0.1.4      2021-04-26 [1] CRAN (R 4.3.3)
##  jsonlite               2.0.0      2025-03-27 [1] CRAN (R 4.3.3)
##  kableExtra           * 1.4.0      2024-01-24 [1] CRAN (R 4.3.3)
##  knitr                * 1.50       2025-03-16 [1] CRAN (R 4.3.3)
##  lattice                0.22-7     2025-04-02 [1] CRAN (R 4.3.3)
##  lifecycle              1.0.4      2023-11-07 [1] CRAN (R 4.3.3)
##  lubridate            * 1.9.4      2024-12-08 [1] CRAN (R 4.3.3)
##  magrittr               2.0.3      2022-03-30 [1] CRAN (R 4.3.3)
##  MASS                   7.3-60.0.1 2024-01-13 [1] CRAN (R 4.3.3)
##  Matrix                 1.6-5      2024-01-11 [1] CRAN (R 4.3.3)
##  MatrixGenerics         1.14.0     2023-10-24 [1] Bioconductor
##  matrixStats            1.5.0      2025-01-07 [1] CRAN (R 4.3.3)
##  mgcv                   1.9-2      2025-04-02 [1] CRAN (R 4.3.3)
##  munsell                0.5.1      2024-04-01 [1] CRAN (R 4.3.3)
##  nlme                   3.1-168    2025-03-31 [1] CRAN (R 4.3.3)
##  patchwork            * 1.3.0      2024-09-16 [1] CRAN (R 4.3.3)
##  permute                0.9-7      2022-01-27 [1] CRAN (R 4.3.3)
##  pillar                 1.10.1     2025-01-07 [1] CRAN (R 4.3.3)
##  pinfsc50               1.3.0      2023-12-05 [1] CRAN (R 4.3.3)
##  pkgconfig              2.0.3      2019-09-22 [1] CRAN (R 4.3.3)
##  plyranges            * 1.22.0     2023-10-24 [1] Bioconductor
##  png                    0.1-8      2022-11-29 [1] CRAN (R 4.3.3)
##  purrr                * 1.0.4      2025-02-05 [1] CRAN (R 4.3.3)
##  R6                     2.6.1      2025-02-15 [1] CRAN (R 4.3.3)
##  Rcpp                   1.0.14     2025-01-12 [1] CRAN (R 4.3.3)
##  RCurl                  1.98-1.16  2024-07-11 [1] CRAN (R 4.3.3)
##  readr                * 2.1.5      2024-01-10 [1] CRAN (R 4.3.3)
##  readxl               * 1.4.5      2025-03-07 [1] CRAN (R 4.3.3)
##  restfulr               0.0.15     2022-06-16 [1] CRAN (R 4.3.3)
##  RIdeogram            * 0.2.2      2020-01-20 [1] CRAN (R 4.3.3)
##  rjson                  0.2.23     2024-09-16 [1] CRAN (R 4.3.3)
##  rlang                  1.1.5      2025-01-17 [1] CRAN (R 4.3.3)
##  rmarkdown              2.29       2024-11-04 [1] CRAN (R 4.3.3)
##  Rsamtools              2.18.0     2023-10-24 [1] Bioconductor
##  rstatix                0.7.2      2023-02-01 [1] CRAN (R 4.3.3)
##  rstudioapi             0.17.1     2024-10-22 [1] CRAN (R 4.3.3)
##  rsvg                   2.6.1      2024-09-20 [1] CRAN (R 4.3.3)
##  rtracklayer            1.62.0     2023-10-24 [1] Bioconductor
##  S4Arrays               1.2.0      2023-10-24 [1] Bioconductor
##  S4Vectors            * 0.40.2     2023-11-23 [1] Bioconductor 3.18 (R 4.3.3)
##  sass                   0.4.9      2024-03-15 [1] CRAN (R 4.3.3)
##  scales               * 1.3.0      2023-11-28 [1] CRAN (R 4.3.3)
##  sessioninfo          * 1.2.3      2025-02-05 [1] CRAN (R 4.3.3)
##  SparseArray            1.2.2      2023-11-07 [1] Bioconductor
##  stringi                1.8.7      2025-03-27 [1] CRAN (R 4.3.3)
##  stringr              * 1.5.1      2023-11-14 [1] CRAN (R 4.3.3)
##  SummarizedExperiment   1.32.0     2023-10-24 [1] Bioconductor
##  svglite                2.1.3      2023-12-08 [1] CRAN (R 4.3.3)
##  systemfonts            1.2.1      2025-01-20 [1] CRAN (R 4.3.3)
##  tibble               * 3.2.1      2023-03-20 [1] CRAN (R 4.3.3)
##  tidyr                * 1.3.1      2024-01-24 [1] CRAN (R 4.3.3)
##  tidyselect             1.2.1      2024-03-11 [1] CRAN (R 4.3.3)
##  tidyverse            * 2.0.0      2023-02-22 [1] CRAN (R 4.3.3)
##  timechange             0.3.0      2024-01-18 [1] CRAN (R 4.3.3)
##  tzdb                   0.5.0      2025-03-15 [1] CRAN (R 4.3.3)
##  vcfR                 * 1.15.0     2023-12-08 [1] CRAN (R 4.3.3)
##  vctrs                  0.6.5      2023-12-01 [1] CRAN (R 4.3.3)
##  vegan                  2.6-10     2025-01-29 [1] CRAN (R 4.3.3)
##  viridis                0.6.5      2024-01-29 [1] CRAN (R 4.3.3)
##  viridisLite            0.4.2      2023-05-02 [1] CRAN (R 4.3.3)
##  vroom                  1.6.5      2023-12-05 [1] CRAN (R 4.3.3)
##  withr                  3.0.2      2024-10-28 [1] CRAN (R 4.3.3)
##  xfun                   0.52       2025-04-02 [1] CRAN (R 4.3.3)
##  XML                    3.99-0.17  2024-06-25 [1] CRAN (R 4.3.3)
##  xml2                   1.3.8      2025-03-14 [1] CRAN (R 4.3.3)
##  XVector                0.42.0     2023-10-24 [1] Bioconductor
##  yaml                 * 2.3.10     2024-07-26 [1] CRAN (R 4.3.3)
##  zlibbioc               1.48.0     2023-10-24 [1] Bioconductor
## 
##  [1] /data/cephfs-1/work/groups/cubi/users/vonkunic_c/.stem-cnv-check/f4f9fb2443eadf59b8a94b12a19c1c0f_/lib/R/library
##  * ── Packages attached to the search path.
## 
## ──────────────────────────────────────────────────────────────────────────────
```

### CNV calling

#### de-novo CNV calls

de-novo CNV calls table

⇩

This section describes all de-novo CNV calls, meaning calls without a
match in the reference sample. The table allows sorting and filtering
the calls by various criteria, default is sorting by Check-Score. The
Check-Score is described on our upcoming manuscript and combines
contributions from CNV size and copynumber as well as additions from
annotation from overlapping
stem
cell hotspots, cancer driver genes, predicted dosage sensitive genes
and other gene annotations.

Hovering over the column headers gives explanations for each column
and the “Column visibility” button can be used to change the default
selection of visible columns.

The section immediately below the table contains details for each CNV
call, including a plot of the CNV region, (if relevant) a table of
annotated genes and hotspots, and a table of all genes overlapping the
CNV (or plot region).

##### nr1-cbs\_dup\_chr2\_47806765\_47900182

##### nr2-combined-call\_loss\_chr22\_33707202\_33849413

##### nr3-combined-call\_loss\_chr13\_32042832\_32157759

##### nr4-cbs\_del\_chr7\_49866430\_49971392

##### nr5-cbs\_dup\_chr19\_47042461\_47065746

##### nr6-cbs\_del\_chr4\_69273240\_69294494

##### nr7-penncnv\_del\_chr11\_25680755\_25743637

##### nr8-cbs\_del\_chr11\_14769567\_14830944

##### nr9-cbs\_dup\_chr12\_7609671\_7651050

##### nr10-cbs\_dup\_chr9\_93126535\_93163919

##### nr11-cbs\_del\_chrx\_131100759\_131140251

```
## No genes in the call area.
```

##### nr12-cbs\_del\_chrx\_45296827\_45334004

##### nr13-cbs\_del\_chrx\_32319433\_32353128

##### nr14-cbs\_del\_chr4\_145387726\_145422391

```
## No genes in the call area.
```

##### nr15-cbs\_del\_chr11\_87974459\_88006065

##### nr16-cbs\_del\_chr2\_153683688\_153713500

```
## No genes in the call area.
```

##### nr17-cbs\_del\_chr7\_6074927\_6103247

##### nr18-combined-call\_loss\_chr5\_17483817\_17509779

##### nr19-penncnv\_dup\_chr3\_127068021\_127087392

```
## No genes in the call area.
```

##### nr20-cbs\_dup\_chr2\_197308812\_197323059

#### Reference genotype CNV calls

reference genotype CNV calls table

⇩

This section describes all reference CNV calls, meaning calls for
which a match in the reference sample was found. Matching of CNV calls
is based on a minimum of at least 50% reciprocal overlap between sample
and reference. Otherwise this section uses the same layout as to the
de-novo calls section.

##### nr1-combined-call\_loss\_chr16\_63870714\_64297219

##### nr2-combined-call\_loss\_chr1\_196743486\_196852356

##### nr3-combined-call\_loss\_chr3\_4069115\_4171652

##### nr4-penncnv\_loh\_chr5\_121811523\_122297372

##### nr5-cbs\_dup\_chr3\_127019606\_127166972

##### nr6-penncnv\_loh\_chr6\_149889403\_150249731

##### nr7-penncnv\_loh\_chr16\_76238719\_76652989

##### nr8-penncnv\_loh\_chr18\_60266934\_60681794

##### nr9-penncnv\_loh\_chr10\_130378643\_130789730

##### nr10-penncnv\_loh\_chr2\_37504234\_37877670

##### nr11-penncnv\_loh\_chr6\_32530203\_32817006

##### nr12-cbs\_dup\_chr10\_133534395\_133620799

##### nr13-penncnv\_loh\_chr10\_133463341\_133620799

##### nr14-penncnv\_loh\_chr6\_31316549\_31459190

##### nr15-cbs\_del\_chr6\_141752404\_141798326

```
## No genes in the call area.
```

##### nr16-combined-call\_loss\_chr13\_37496412\_37540475

##### nr17-cbs\_del\_chr11\_7793392\_7825582

##### nr18-combined-call\_loss\_chr10\_55204437\_55228596

##### nr19-cbs\_del\_chr11\_25680755\_25705515

```
## No genes in the call area.
```

##### nr20-penncnv\_del\_chr6\_141752404\_141774564

```
## No genes in the call area.
```

### SNV analysis

#### Table of de-novo SNVs

SNV table explanations

⇩

This table lists all SNVs detected by the Chip Array which are
different from the reference genome and are annotated as at least
protein changing. Due to their potential impact these are now called
“SNVs” rather than “SNPs”, independent of their actual (unknown)
frequency in the population.

All SNVs are categorised into one of the following categories (shown
in the hidden SNV category column):

- hotspot-match: SNV matching a known stemcell hotspot mutation
  (see also SNV hotspot coverage)
- hotspot-gene: SNV in a gene with known iPSC hotspots (see also
  SNV hotspot coverage)
- protein-ablation: SNV (likely) fully disrupting protein function
  (i.e. frameshift, stop gain, stop loss)
- protein-changing: SNV causing a change the protein sequence
  (i.e. missense, inframe)
- other: SNV with other unclear or undetermined effect on protein
  function

The “SNV label” further categorizes the SNVs into:

- Critical de-novo: SNV with likely critical significance on hiPSC
  line
- Reportable de-novo: SNV with possible significance on hiPSC
  line
- Unreliable critical/reportable: SNV with likely or possible
  significance on hiPSC line, but unreliable signal
- de-novo SNV: SNV with de-novo status, but no clear functional
  impact
- Reference genotype: SNV already detected in the reference
  sample

The following categories are assinged as “Critical” or “Reportable”
(de-novo):

- Critical de-novo: hotspot-match
- Reportable de-novo: hotspot-gene
- Reportable de-novo: protein-ablation

A complete, up-to-date list of all stem cell SNV hotspots is also
available
online.

The table allows sorting and filtering the SNVs by various criteria,
default is sorting by the `SNV Label`.Hovering over the
column headers gives explanations for each column and the “Column
visibility” button can beused to show (or hide) columns. Each SNV
genotype (GT) is shown in vcf format: each allele is represented by a
single number, separated by a forward slash. A 0 indicates the reference
allele, a 1 indicates the alternate allele. A dot (.) indicates that the
genotype could not be determined.

#### Table of reference SNVs

#### SNV hotspot coverage

SNV hotspot coverage explanations

⇩

This table lists all genes that have known point mutation hotspots
for stem cells, a source for the hotspots, the selected primary
transcript for each gene, as well as the coverage of the genes on cDNA,
CDS and protein level (percent coverage of bases/amino acids, as well as
absolute numbers). The coverage is based on all probes contained on the
utilised array.

The “Hotspots” column, lists the specific annotated protein changes for
each gene and whether or not any probe on the array covers each of the
specific mutations.

#### SNV QC details

### Sample comparison

#### Genome Overview

Genome overview explanations

⇩

The following plots each show a whole chromosome overview of the
sample, combining to a whole genome view. CNV calls (filtered based on
the config settings) are shown as colored background bars, with the
color indicating the type of call: green for gains, red for losses, and
grey for LOH. Additionally, if the sample has a reference, SNVs that are
labelled critical or protein changing/unreliable critical are also
highlighted in red and orange, respectively.

##### chr1

##### chr2

##### chr3

##### chr4

##### chr5

##### chr6

##### chr7

##### chr8

##### chr9

##### chr10

##### chr11

##### chr12

##### chr13

##### chr14

##### chr15

##### chr16

##### chr17

##### chr18

##### chr19

##### chr20

##### chr21

##### chr22

##### chrX

##### chrY

#### Identity comparison

Dendrogram explanations

⇩

Sample identities can be comparsed based on the dendrogram built on
the SNP genotypes. The dendrogram is built using the manhattan distance
between samples, counting both alleles from Probes that are not quality
in every included sample. Accordingly, the distance between two samples
is the sum of the absolute differences between the two alleles at each
SNP (also shown in the table below) after QC filters. Samples that are
very close together are likely identical or clonally related. Sample
selection as well as color and shape lables are controlled by the config
file.

Only 15 shapes are available, but “Chip\_Name” would need 17. Consider
using fewer unqiue entries. These values are summarised as “Other”:
208305080104, 209362520148
